# Development of fluorescence polarization-based competition assay for nicotinamide *N*-methyltransferase

**DOI:** 10.1101/2020.04.10.034793

**Authors:** Iredia D. Iyamu, Rong Huang

## Abstract

Methylation-mediated pathways play an important role in the progression of cancer. Inhibitors of several key methyltransferases including DNA methyltransferases and histone methyltransferases have proven to be instrumental for both understanding the function of the respective enzymes activites and translational applications in cancer epigenetic therapy. Nicotinamide *N*-methyltransferase (NNMT) is a major metabolic enzyme involved in epigenetic regulation through catalysis of methyl transfer from the cofactor *S*-adenosyl-L-methionine, onto nicotinamide and other pyridines, to form S-adenosyl homocysteine and 1-methyl-nicotinamide or the corresponding pyridinium ions. Accumulating evidence infers that NNMT is a novel therapeutic target for a variety of diseases such as cancer, diabetes, obesity, cardiovascular and neurodegenerative diseases. Therefore, there is an urgent need to discover potent and specific inhibitors for NNMT. Herein, we reported the design and synthesis of a fluorescent probe **II138**, and established a fluorescence polarization (FP)-based competition assay for evaluation of NNMT inhibitors. Importantly, the unique feature of this FP competition assay is its capability to identify inhibitors that interfere with the interaction of the NNMT active site directly or allosterically. In addition, this assay performane is robust with a Z factor of 0.76, and applicable in high-throughput screening for inhibitors for NNMT.

## Introduction

Nicotinamide N-methyltransferase (NNMT) is a metabolic enzyme that catalyzes the transfer of a methyl group from the cofactor S-adenosyl-L-methionine (AdoMet) to nicotinamide (NAM) and other pyridines to produce S-adenosyl-L-homocysteine (AdoHcy) and respective methylated products. NNMT is expressed predominantly in the liver and adipose tissue, and at lower levels in other organs [1]. Through catalyzing N-methylation of pyridine-containing compounds, NNMT plays an important role in the biotransformation and detoxification of many drugs and xenobiotic compounds in the human body. NAM, a form of vitamin B3, is the most studied endogenous substrate of NNMT and is converted to N1-methylnicotinamide (MNAM) after methylation. NAM is also a precursor of salvage pathways to produce NAD^+^, an important coenzyme involved in redox metabolism and sirtuins-catalyzed deacetylation reaction [2,3]. In addition, NNMT also methylates 4-phenylpyridine to generate the neurotoxin 1-methyl-4-phenylpyridinium ion, which interferes with complex I and is implicated in idiopathic Parkinson’s disease [4].

The dual functions of NNMT in the metabolism of both NAM and AdoMet manifest its significance in cellular metabolism and epigenetic pathways. Not surprisingly, upregulation of NNMT has been implicated in cancers, obesity, diabetes, cardiovascular and neurodegenerative diseases [3–10]. Genetic studies have shown that the loss of NNMT decreases cell proliferation, migration, and/or metastasis of various cancer cells [11]. In addition, knockdown of NNMT has led to an increase of metabolism and prevention of weight gain [3]. The emerging importance of NNMT imposes the need to develop selective and cell-potent inhibitors to elucidate its therapeutic potential. Despite recent progresses in the development of NNMT bisubstrate inhibitors achieving inhibitory activity at nM levels, limited cell permeability restrains their applications in cellular studies [12–14]. Meanwhile, available small molecule inhibitors mainly only bear moderate activity at µM levels [15,16]. Hence, there remains an urgent need for new cell-potent inhibitors to delineate the physiological and pathological functions of NNMT.

To facilitate the discovery of NNMT inhibitors, it is important to have a convenient and efficient assay for NNMT to screen and evaluate the inhibitors. Currently, a widely used assay is an enzymatic assay to indirectly monitor the production of AdoHcy, either through an AdoHcy hydrolase (SAHH)-coupled fluorescence or a bioluminescent MTase-Glo™ assay (Promega, Inc). Both assays require additional coupling enzymes and reagents to produce a detectable signal. In the SAHH-coupled fluorescence assay, AdoHcy is hydrolyzed by SAHH to generate homocysteine, which is then reacted with a thiol-specific dye to form a strong fluorescent signal [17,18]. The MTase-Glo™ assay needs several coupling enzymes to convert AdoHcy to ADP and produce a luciferase signal [17,19]. Direct assay to measure the methylation product like MNAM has been developed through the use of high-pressure liquid chromatography (HPLC) [20]. However, HPLC is time-consuming and difficult to apply to high throughput screening (HTS) applications. Additionally, non-coupled assays that use quinoline as a NNMT substrate enable a direct detection of the fluorescence of the corresponding methylation product methylquinolinium at 405 nm, but this assay has a low signal-to-noise ratio [21,22]. In addition, biophysical methods like isothermal titration calorimetry and surface plasmon resonance are useful techniques for direct determination of binding affinities of inhibitors to the target. However, the requirements of expensive instruments and high stability of the protein have limited their applications as a primary method for screening inhibitors. On the other hand, fluorescence polarization (FP) is a HTS-amenable method to characterize the binding of a fluorescent ligand to the target or any compound that interferes with the aforementioned interaction in solution, making it very attractive [23]. In this work, we report the design and synthesis of the first fluorescent probe **II138** for NNMT to our knowledge. Furthermore, we have established and optimized a FP-based competition assay in an HTS-compatible format, which can be easily applied for identification and evaluation of NNMT inhibitors that perturb the binding to the active site either directly or allosterically.

## 2. Materials and methods

### 2.1 Materials and instruments

The reagents and solvents were purchased from commercial sources (Fisher and Sigma-Aldrich) and used directly. Analytical thin-layer chromatography was performed on ready-to-use plates with silica gel 60 (Merck, F254). Flash column chromatography was performed over silica gel (grade 60, 230−400 mesh) on Teledyne Isco CombiFlash purification system. Final compounds were purified on preparative reversed phase high-pressure liquid chromatography (RP-HPLC) and was performed on Agilent 1260 Series system with Agilent 1260 Infinity II Variable Wavelength Detector (G7114A, UV = 254 nM) and a Waters BEH C18 (130Å, 5 *µ*m, 10 mm X 250 mm) column. Final compounds were eluted using a gradient 95% water (with 0.1% formic acid) and 5% acetonitrile (with 0.1% formic acid) at a flow rate of 4 mL/min over 30 min.

NMR spectra were acquired on a Bruker AV500 instrument (500 MHz for 1H-NMR, 126 MHz for 13C-NMR). TLC-MS were acquired using Advion CMS-L MS. The Agilent 1260 Infinity II Variable Wavelength Detector (G7114A, UV = 254 nM) and an Agilent ZORBAX RR SB-C18 (80Å, 3.5 *µ*m, 4.6 x 150 mm) at a flow rate of 1 mL/min using a solvent system of 100% water with 0.1% TFA to 40 or 60% methanol over 20min were used to assess purity of final compounds. All the purity of target compounds showed >95% in RP-HPLC. FP was monitored on a BMG CLARIOstar microplate reader.

### 2.2 Protein Expression and Purification

Expression and purification of full-length human NNMT wild type was performed as previously described [24]. Briefly, full-length hNNMT (amino acids 1-270) was codon optimized, synthesized, and cloned into pET28a(+)-TEV vector (GenScript). Protein was expressed in *E. coli* BL21-CodonPlus(DE3)-RIPL competent cells in LB media with kanamycin and induced by 0.3 mM isopropyl-D-1-thiogalactopyranoside at 16 °C for 20 hours. Harvested cells were lysed through sonication (Qsonica Q55 cell disruptor) on ice in 10 volumes of 50 mM KH_2_PO_4_/K_2_HPO_4_ (pH = 7.4) containing 500 mM NaCl, 25 mM imidazole, 5 mM ß-mercaptoethanol, and 100 µM PMSF. The cell lysate was centrifuged at 25,000 × g for 30 minutes at 4 °C. Then the supernatant was loaded to the HiTrap FF Ni-NTA column on a GE AKTA Prime purification system and eluted with a step gradient of imidazole (0 to 0.5 M), 50 mM KH_2_PO_4_/K_2_HPO_4_ (pH = 7.4), 500 mM NaCl, and 0.5 mM TCEP. The peak fractions were verified by SDS-PAGE analysis, combined to dialyze in the dialysis buffer (25 mM Tris, pH = 7.5, 150 mM NaCl, 50 mM KCl), and concentrated to 1.5 mg/mL.

### 2.3 FP measurement, binding and competition assay

All FP measurements were performed on a BMG CLARIOstar microplate reader in black opaque 384-well microplates (Corning #3820) with excitation 482 nm and emission 530 nm. All experiments were performed in triplicates in a volume of 20 µL per well in 25 mM Tris, 50 mM KCl, 0.01% Tween pH 7.5. For direct binding assay, different concentrations of NNMT were titrated against the probe at the concentration of 5 nM. The polarization (mP) was measured after both 30 min and 60 min incubation at room temperature. The dissociation constant (*K*_*d*_^*app*^) was obtained by fitting the fluorescence polarization values and the corresponding protein concentrations into a nonlinear regression model in GraphPad Prism 8. For the competition binding assay, a solution of the enzyme (NNMT) at 0.5 µM concentration was incubated with different concentrations of the inhibitor at room temperature for 30 mins. FP probe was then added to yield a final concentration of 5 nM. The polarization was measured after the mixture was incubated for 30 mins. The FP values were plotted against the log of inhibitor concentrations into a nonlinear regression model. The *K*_*i*_ were calculated from Binding-Competitive model in GraphPad Prism [25].

### 2.4 Z’-factor determination and data analysis

The fluorescence polarization signals of 100 wells of negative control (no inhibitors added to the competition assay) and 100 wells of positive control samples (5 µM of bisubstrate inhibitor **LL320**, Figure 1) were measured in a single 384-well plate after 60 mins incubation at room temperature. The statistical parameter Z-factor was calculated based on the following equation:

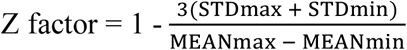

**Figure 1.**
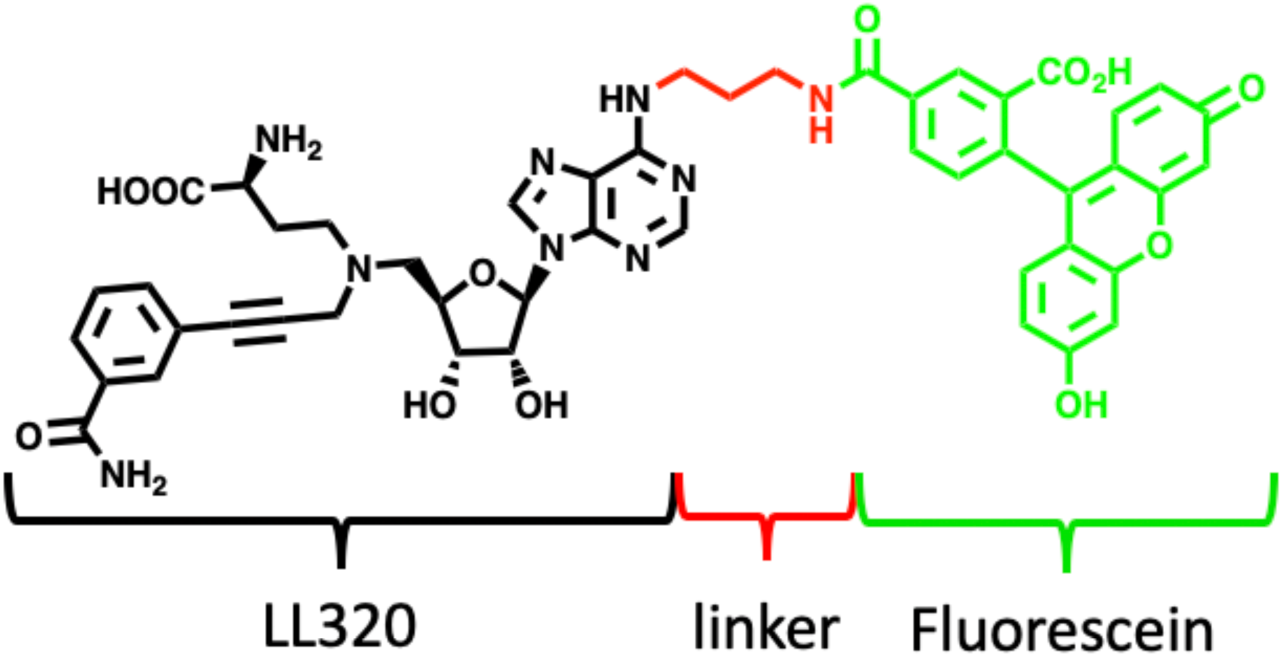
Structure of FP probe **II138** comprised of **LL320** (black), the linker (red), and fluorescein (green).

## 3. Results and Discussion

### Probe Design

The inherent feature of the NNMT active site imposes a challenge on the discovery of a potent and specific inhibitor by solely targeting NAM or AdoMet binding site because of the relatively small binding site of NAM and conserved binding site of AdoMet. We have recently reported the development of a potent and selective propargyl-linked bisubstrate inhibitor **LL320** (*K*_i_ = 1.6 ± 0.3 nM) for NNMT, displaying over 1,000-fold selectivity for a panel of MTases [26]. Inspired by the high potency and selectivity of **LL320**, we hypothesize that a fluorescent derivative of **LL320** would enable the development of a FP assay for the identification of potent and specific NNMT inhibitors. Because **LL320** occupies both NAM and AdoMet binding sites, the advantage of this FP assay is to allow the discovery of NNMT inhibitors through various mechanisms, such as directly or allosterically binding to either NAM or AdoMet binding site, or both. Our designed FP probe (**II138**) is comprised of three parts: **LL320**, linker, and the fluorescent dye (Figure 1). We chose fluorescein as the dye because of its high quantum yield and availability. The position, flexibility and size of the linker of the FP probe is crucial in order to retain its binding affinity for NNMT. Examination of the co-crystal structure of NNMT-**LL320** (PDB ID: 6PVS) suggests that the *N*^*6*^ of adenosine is a favorable site to attach a linker because it is exposed towards the solvent (Figure 2A). In addition, the distance between the *N*^*6*^ and the surface of NNMT is about 4.2 Å (Figure 2B). Hence, we hypothesize that a 3-carbon linker will be enough to link LL320 to a fluorescent dye while minimizing the adverse effect of the fluorescein on the binding affinity for NNMT. To allow certain flexibility of the linker, a propyl group was designed to connect the fluorescein with **LL320** to produce **II138** (Figure 1).

**Figure 2.**
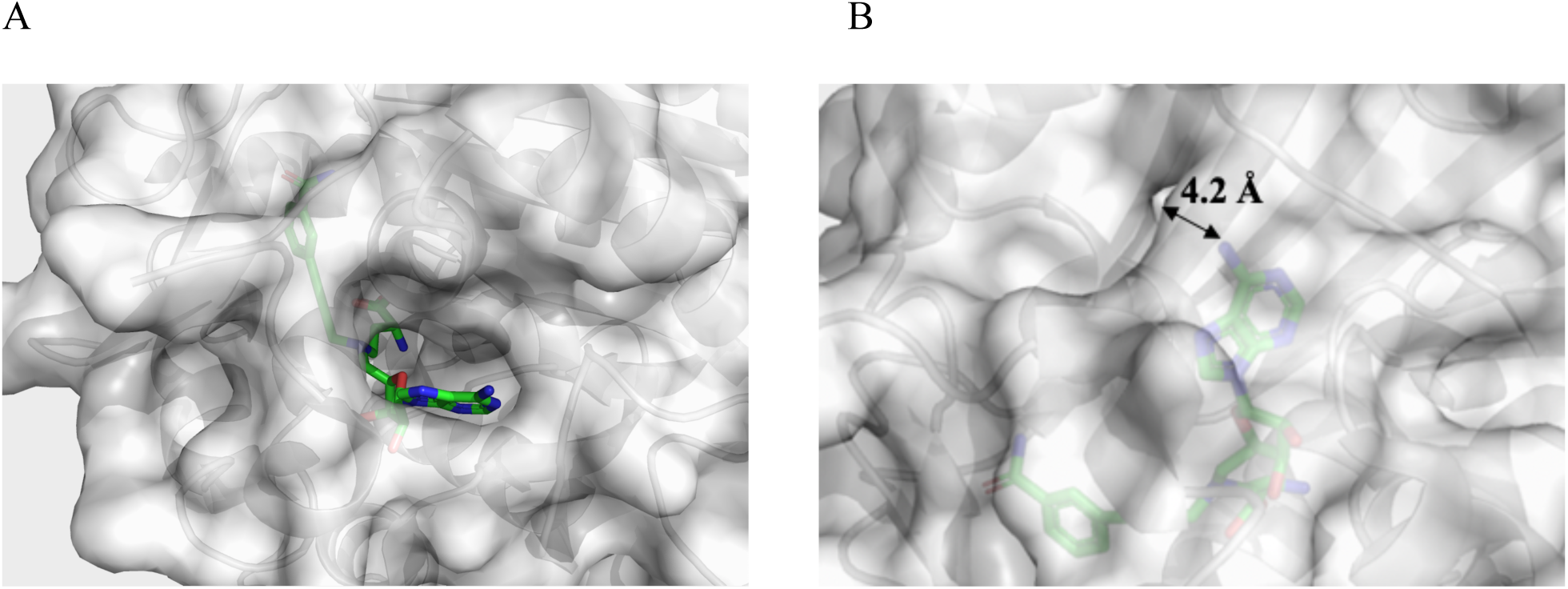
Crystal structure of hNNMT-LL320 complex (PDB: 6PVS). A. hNNMT is colored gray and **LL320** is shown in green stick with *N*^*6*^ adenosine pointing towards the solvent. B. The distance between *N*^*6*^ and the protein surface is 4.2 Å.

### Synthesis of FP probe

The FP probe was synthesized according to Scheme 1. First, the hydroxyl group in commercially available (2*R*,3*R*,4*S*,5*R*)-2-(6-chloro-9*H*-purin-9-yl)-5-(hydroxymethyl)tetrahydrofuran-3,4-diol **1** was protected by treatment with *p*-toluenesulfonic acid in acetone to afford the acetal **2**. Then, the amino linker at *N*^*6*^ was incorporated by substitution with propane-1,3-diamine which was subsequently protected upon treatment with ethyl trifluoroacetate to yield **3** [27]. The subsequent Mitsunobu reaction with isoindoline-1,3-dione afforded the masked amine **4**, which was deprotected by hydrazine to produce **5**. Reductive amination of **5** with *tert*-butyl (*S*)-2-((*tert*-butoxycarbonyl)amino)-4-oxobutanoate afforded **6**, which was subjected to a second reductive amination with 3-(3-oxoprop-1-yn-1-yl)benzonitrile to yield **7**. Oxidization of the cyano group of **7** to a primary amide by hydrogen peroxide followed by removal of the TFA ester provided **9**. Simultaneous removal of the acetal, boc and *tert*-butoxyl protecting groups was achieved by TFA treatment to yield **10**, which was used to investigate the effect of the propane-1,3-diamine linker on **LL320**. To synthesize the FP probe **II138**, compound **9** was subjected to the amidation reaction with 5-carboxyfluorescein to attach the fluorescein followed by global deprotection (Scheme 1).

**Scheme 1.**
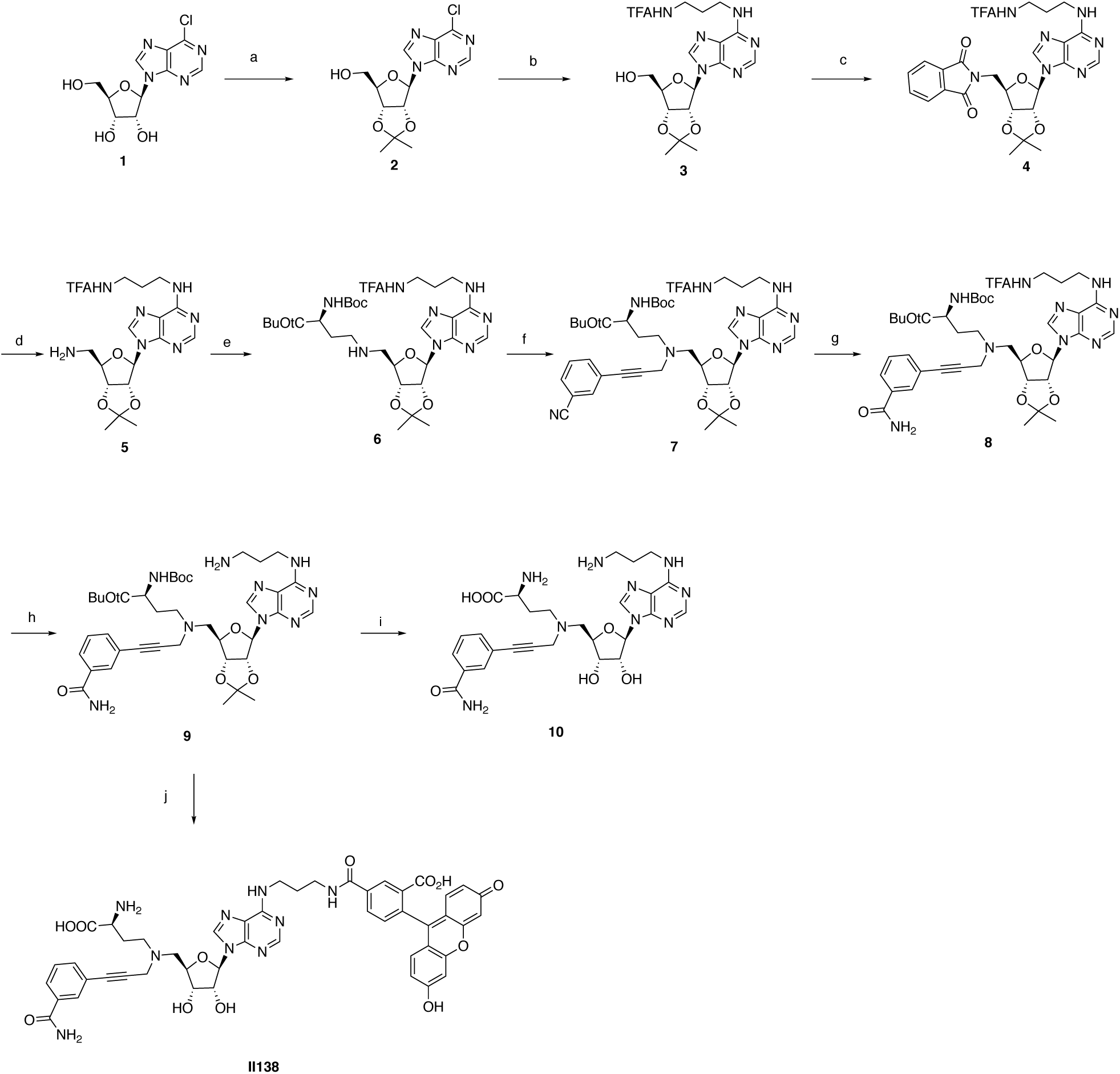
Synthesis of FP probe^a^. ^a^Reagents and conditions: (a) *p*TsOH, acetone, 3 h, quant. yield; (b) propane-1,3-diamine, TEA, EtOH, 60 oC, 2 h then; CF_3_CO_2_Et, TEA, MeOH, 12 h, 89 %; (c) isoindoline-1,3-dione, PPh_3_, DIAD, THF, 2 h, 90 %; (d) NH_2_NH_2_, MeOH, 8 h, 63 %; (e) *tert*-butyl (*S*)-2-((*tert*-butoxycarbonyl)amino)-4-oxobutanoate, NaBH_3_CN, AcOH, MeOH, 2 h, 67 %; (f) 3-(3-oxoprop-1-yn-1-yl)benzonitrile, NaBH_3_CN, AcOH, MeOH, 2 h, 79 %; (g) K_2_CO_3_, H_2_O_2_, DMSO, quant. yield; (h) NH_4_OH, MeOH, 77 %; (i) TFA, H_2_O, 8 h, 37 %; (j) fluorescein, DIC, HOBt, DIPEA, DMF; 12 h then TFA, H_2_O, 8 h, 29 %.

### Evaluation of the linker effect

In order to determine the effect of the linker, the *K*_*i*_ of compound **10** was determined by the SAHH-coupled fluorescence assay to be 0.21 ± 0.2 µM, reflecting about 100-fold loss in comparison with **LL320** [26]. This is probably due to the loss of one hydrogen bond donor of the *N*^*6*^ amine in adenosine. Nevertheless, this decreased *K*_*i*_ value is acceptable for the FP probe that is intended for the FP assay development.

### Assay optimization

Both HEPES and Tris buffer were investigated to evaluate their effects on the sensitivity (detection limit) in this assay. In Tris buffer, 5 nM concentration of the probe showed over 10-fold higher total fluorescence intensity compared to the blank, but in HEPES buffer, a probe concentration of >40 nM was required to display 10-fold higher total fluorescence over the blank. (Figure 3A,3B). In order to determine the linearity and the optimal concentration of the probe (**II138**), the fluorescence intensity of the probe was measured at various concentrations ranging from 0 to 1 µM in the Tris Buffer. We chose the concentration of the probe at 5 nM because it gave about 10-fold higher intensity than the blank with no probe (Figure 3A). The direct binding of the FP probe **II138** to the enzyme NNMT was determined by measuring the change in polarization (ΔmP) in presence of different protein concentration. From the sigmoidal binding curve (Figure 3C), 0.5 µM of NNMT was chosen for the subsequent FP assay as it produces over 50% increase in FP between the free ligand. The *K*_*d*_ value of the probe was determined to be 369 ± 14.4 nM.

**Figure 3.**
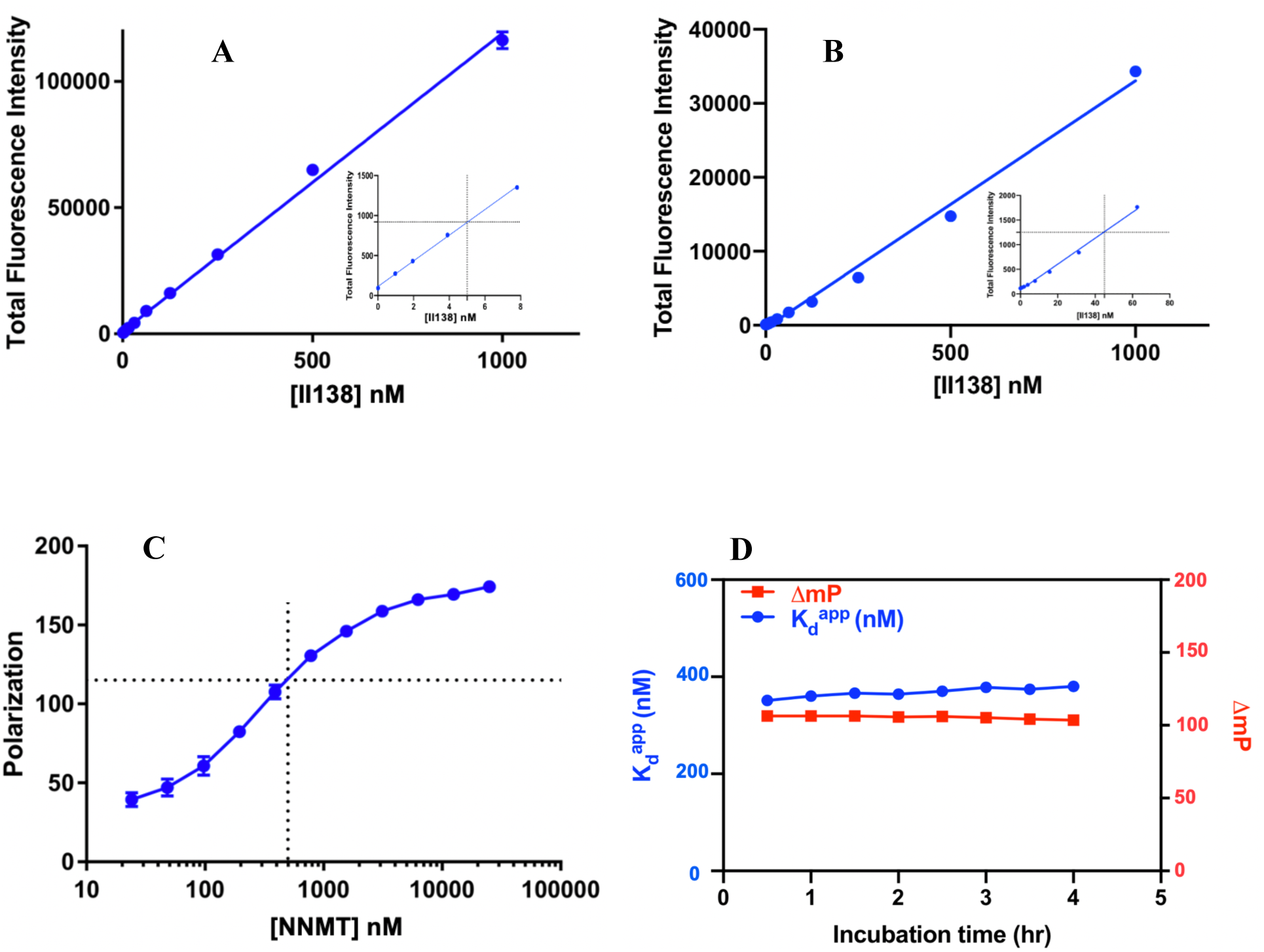
Optimization of FP assay condition. (A) Linearity of the total fluorescence intensity of the probe with increase in probe concentration after incubation without NNMT in Tris buffer. 10-fold increase from blank is zoomed in for clarity. (B) Linearity of the total fluorescence intensity of FP probe after incubation without NNMT in HEPES buffer. (C) Direct Binding of the probe to NNMT with 0.5 µM NNMT concentration indication over 50% polarization signal. (D) Effect of different incubation time on the binding affinity of probe to NNMT.

To evaluate the stability of the probe and the optimal incubation time for the FP binding assay, the reaction mixture was incubated and analyzed at room temperature every 30 minutes for 4 hours. The system reached equilibrium within 30 min and the binding affinity remained stable for the entire 4 h duration of the experiment, indicating that NNMT was stable for the entire incubation time. The assay window was largely unchanged during the entire 4 h period of incubation (Figure 3D). At higher temperatures (30 °C and 37 °C), there was more variability in the binding affinity and assay window, indicating poor stability at higher temperatures. Thus, we proceeded with the measurement after incubating with the probe for 30 minutes at room temperature because the *K*_*d*_ reached equilibrium and remained stable.

### Competition FP assay

After confirming the direct binding of the FP probe to NNMT, a competition FP assay was performed to evaluate the displacement of the bound probe with reported inhibitors. Compound **10**, the unlabeled analog of the FP probe, was employed in a “gold standard” assay. The *K*_*i*_ of the competitor (**10**) was found to be 0.31 ± 0.7 µM from the FP assay (Figure 4), which was comparable to the *K*_*i*_ value of 0.21 ± 0.2 µM determined by the SAHH-coupled fluorescence assay. The difference in both *K*_*i*_ values could be due to the different assay conditions, particularly the enzyme concentration [25]. However, the comparable values establish the applicability of the FP probe to evaluate NNMT inhibitors.

**Figure 4.**
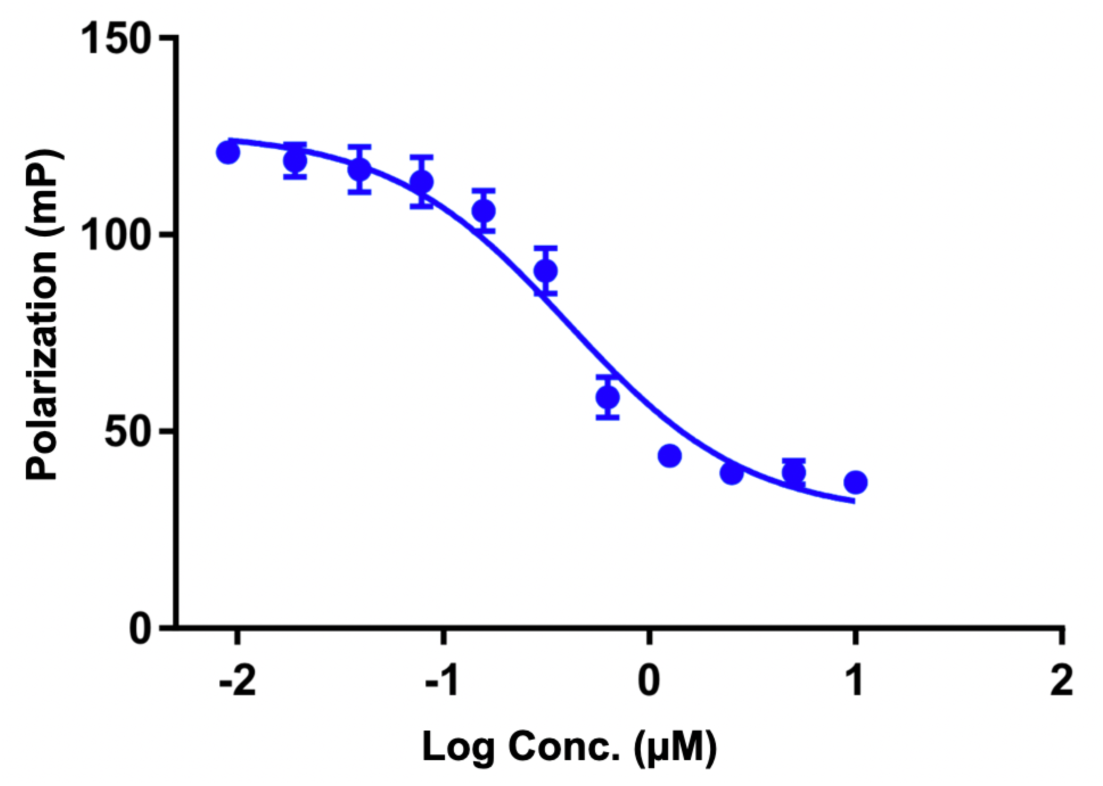
Gold Standard Assay. Competitive Binding assay of the unlabeled probe (**10**) with the FP probe.

### 3.6 Assay performance assessment

In order to further validate potential application of this FP assay, we decided to determine the *K*_*i*_ values of previously reported NNMT inhibitors (Table 1). In general, most inhibitors that displayed a submicromolar to millimolar inhibitory activity showed comparable *K*_*i*_ values in our FP assay in comparison with literature values from other assays. For example, the *K*_*i*_ from our FP assay for AdoHcy was 82.96 ± 14.3 µM, while its IC_50_ was reported to be 75.4 ± 6.3 µM from the UHPLC-MS assay [28]. Meanwhile, two previously-reported bisubstrate inhibitors **VH45** and **MS2756** displayed a *K*_*i*_ value of 59.19 ± 8.1 µM and 11.27 ± 1.8 µM in FP assay versus 29.2 ± 4.0 µM and 10 ± 0.35 µM in the SAHH coupled assay, respectively [13]. Likewise, **LL335** exhibited a *K*_*i*_ value of 5.6 ± 1.2 µM in the FP assay and 3.0 ± 0.83 µM in the SAHH-coupled assay, respectively [26]. However, there is a discrepancy for tight-binding inhibitors that have much higher affinity compared to the binding affinity of the probe **II138**. For instance, **LL319** and **LL320** demonstrated a *K*_*i*_ value of 0.41 ± 0.05 µM and 0.22 ± 0.04 µM in the FP assay, respectively. There is nearly 5- and 130-fold difference for **LL319** and **LL320** in comparison with the values in the SAHH coupled assay, respectively [26]. The difference is resulted from the sensitivity limitation imposed by the binding affinity (*K*_*d*_ =369 ± 14.4 nM) of the FP probe **II138**, which determines the approximate lower end of the inhibitor *K*_*i*_ range measureable by the FP competition assay [29]. Hence, this FP assay will not be effective in distinguishing tight-binding inhibitors, which can usually be resolved by the follow-up activity assay. In addition, we also checked the relative binding of the cofactor and substrate for NNMT. The cofactor AdoMet (*K*_*m*_ of 24 ± 6.8 µM) was determined to have a *K*_*i*_ of 20.27 ± 2.9µM [17]. The substrates NAM and 1-methyl quinoline (1MQ) exhibited over 1 mM *K*_*i*_ in our FP assay, which were in alignment with reported values.

**Table 1.**
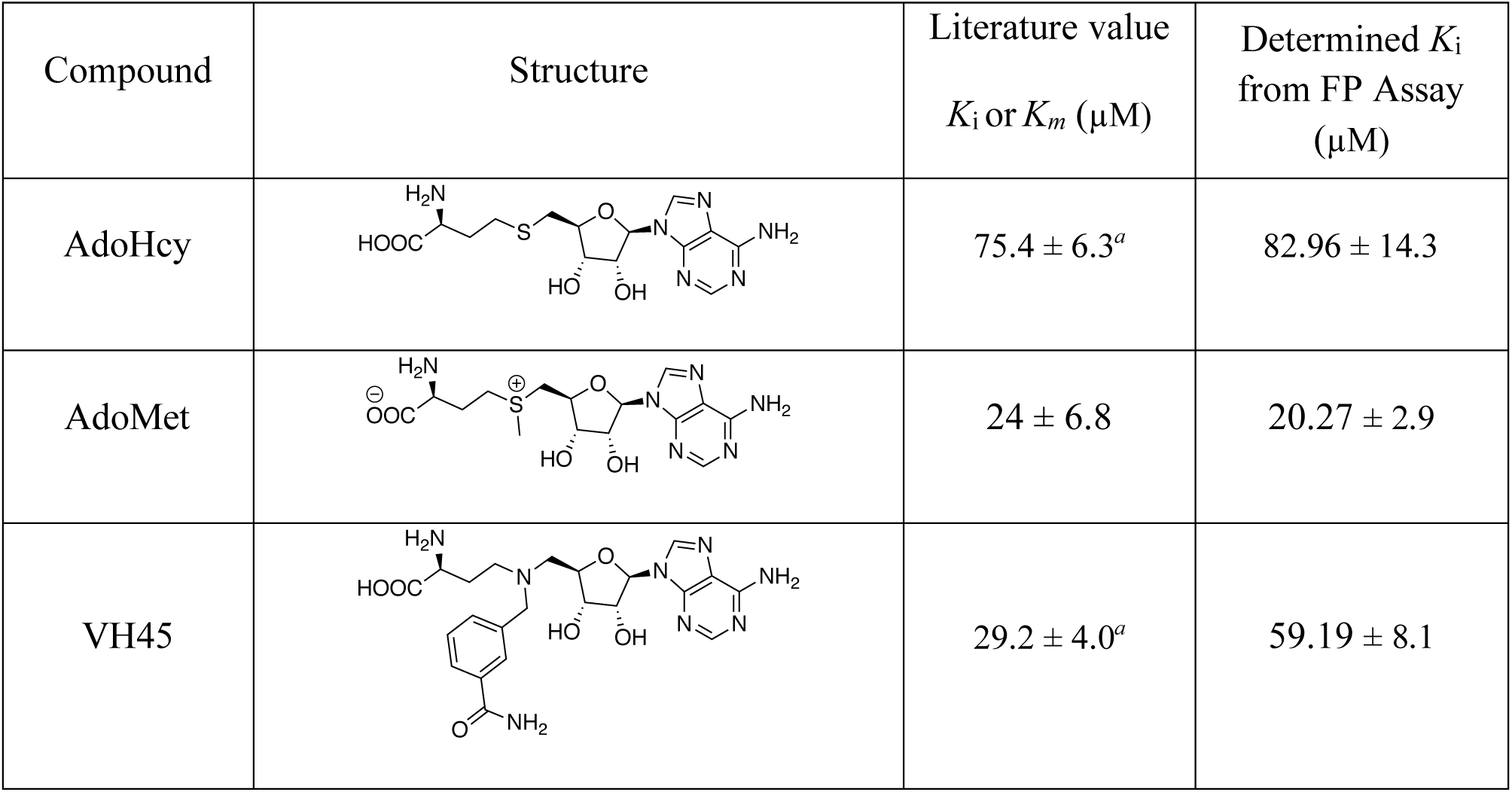
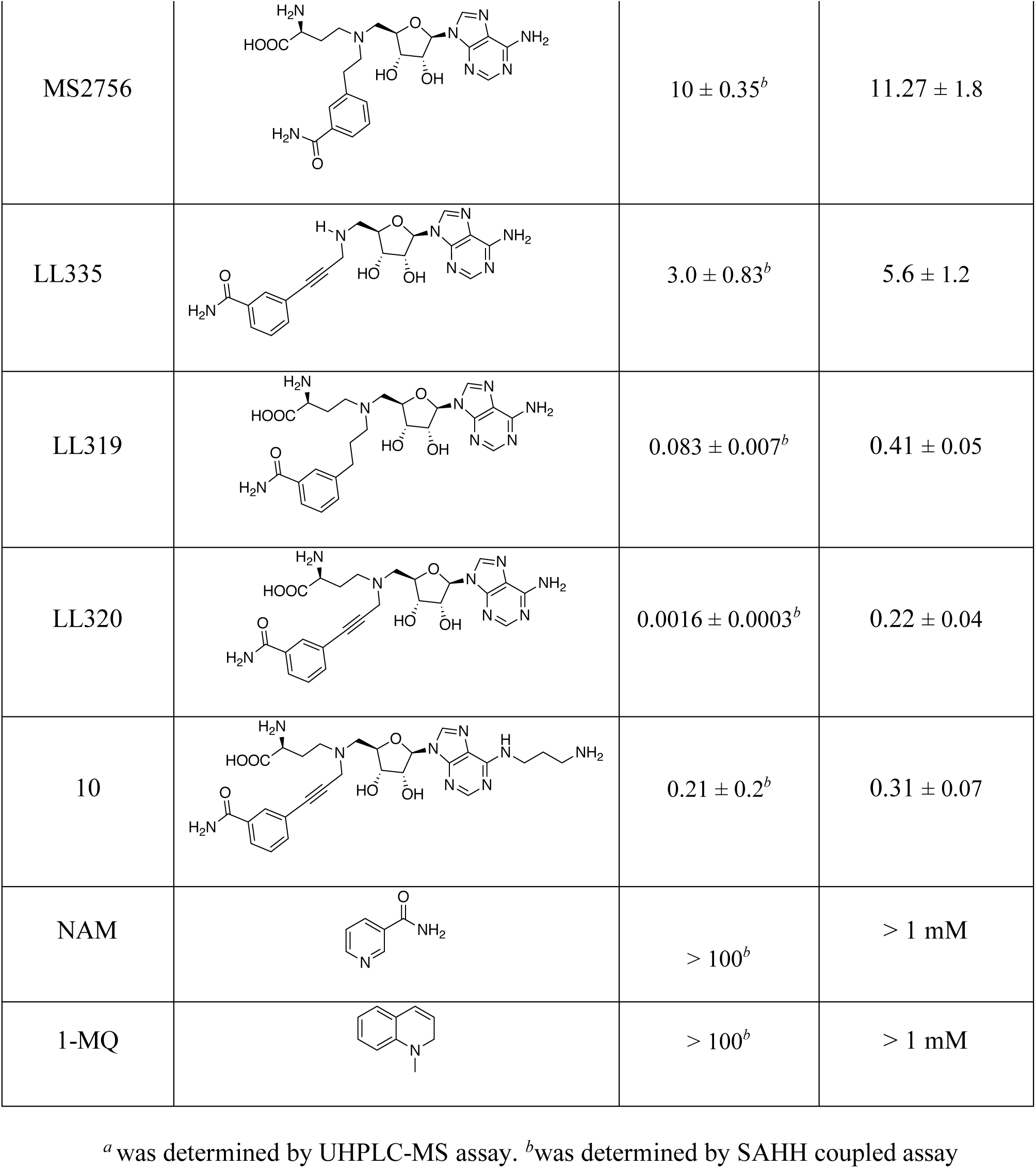
Comparison of *K*_*i*_ values from FP with literature values from other assays

<sup>a</sup> was determined by UHPLC-MS assay. <sup>b</sup> was determined by SAHH coupled assay
| Compound | Structure | Literature value<br>$K_i$ or $K_m$ (μM) | Determined $K_i$<br>from FP Assay<br>(μM) |
| --- | --- | --- | --- |
| AdoHcy | | $75.4 \pm 6.3^a$ | $82.96 \pm 14.3$ |
| AdoMet | | $24 \pm 6.8$ | $20.27 \pm 2.9$ |
| VH45 | | $29.2 \pm 4.0^a$ | $59.19 \pm 8.1$ |
| MS2756 | | $10 \pm 0.35^b$ | $11.27 \pm 1.8$ |
| LL335 | | $3.0 \pm 0.83^b$ | $5.6 \pm 1.2$ |
| LL319 | | $0.083 \pm 0.007^b$ | $0.41 \pm 0.05$ |
| LL320 | | $0.0016 \pm 0.0003^b$ | $0.22 \pm 0.04$ |
| 10 | | $0.21 \pm 0.2^b$ | $0.31 \pm 0.07$ |
| NAM | | $> 100^b$ | $> 1 \text{ mM}$ |
| 1-MQ | | $> 100^b$ | $> 1 \text{ mM}$ |

### Optimization of DMSO compatibility and Z-factor

To evaluate the potential of this probe to be applied for HTS, we proceeded to investigate the effects of DMSO on the FP assay because DMSO is a commonly used solvent to prepare stock solutions for small molecules. The *K*_*d*_ value of the FP probe was examined in the presence of DMSO up to 10% (Figure 5A). The effects of DMSO on the binding affinity of the probe were negligible until DMSO reached 2.5% in the solution. The *K*_*d*_ value remained relatively stable over the 0 to 2.5% DMSO concentration range, with values of 0.34 µM at 0.5% DMSO to 0.38 µM at 2.5% DMSO. Then *K*_*d*_ increased proportionally to the percentage of DMSO from 5%-10%. The assay window (ΔmP) remained relatively stable in the presence of DMSO up to 5%, but it gradually decreased from 5% to 10% DMSO. This may be likely due to the instability of NNMT in the presence of DMSO. Thus, the stability of the *K*_*d*_ and ΔmP in the presence of DMSO up to 2.5% makes this FP assay amenable for high-throughput screening of small molecules with solubility problems.

**Figure 5.**
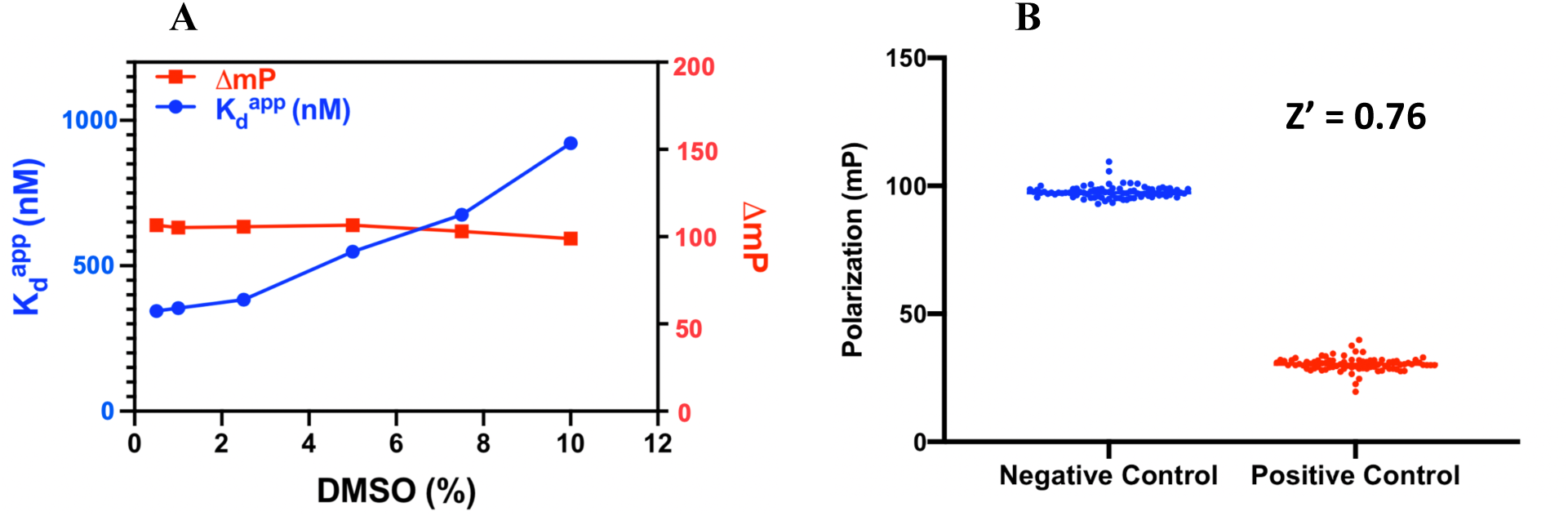
DMSO tolerance and Z’ factor determination (A) Effect of different DMSO concentrations on the binding affinity of probe to NNMT. (B) Z’ factor determination from blue dots (100 negative controls); red dots (100 positive controls).

The statistical parameter Z-factor (Z’) has been used to evaluate the quality of a HTS. Typically, a Z’ value of greater than 0.5 is considered to be an excellent assay. The invariance of the positive and negative control groups resulted in low standard deviation (Figure 5B). The Z’ factor of our developed FP assay was calculated to be 0.76, indicating a robust assay for its application in HTS.

## Conclusion

In summary, we have designed and synthesized a FP probe **II138** for a FP-based competition assay for evaluation of NNMT inhibitors. The probe was determined to directly bind to NNMT with a *K*_d_^app^ value of 369 ± 14.4 nM, which has been used to establish and optimize a FP-based competition assay for measuring the binding affinities of NNMT inhibitors. The requirement for small amount of sample makes this FP competition assay not only an economical but valuable tool for identification and evaluation of NNMT inhibitors. Because of the unique feature of **II138** to occupy both substrate and cofactor binding sites of NNMT, this FP competition assay enables the detection of any molecules that affect the interaction of either the substrate, cofactor, or both with NNMT. Notably, this effect can be either direct or indirect. Moreover, we also demonstrate the robustness of this assay with a Z’value of 0.76 and compatibility with DMSO, indicating that this FP competition assay can be employed in HTS to expedite the discovery of novel NNMT inhibitors.

## EXPERIMENTAL SECTION

### Chemistry General Procedures

The reagents and solvents were purchased from commercial sources (Fisher and Sigma-Aldrich) and used directly. Analytical thin-layer chromatography was performed on ready-to-use plates with silica gel 60 (Merck, F254). Flash column chromatography was performed over silica gel (grade 60, 230−400 mesh) on Teledyne Isco CombiFlash purification system. Final compounds were purified on preparative reversed phase high-pressure liquid chromatography (RP-HPLC) was performed on Agilent 1260 Series system. Systems were run with 0-50% methanol/water gradient with 0.1% TFA. NMR spectra were acquired on a Bruker AV500 instrument (500 MHz for ^1^H-NMR, 126 MHz for ^13^C-NMR).

### General procedure A (Reductive amination)

To a solution of NAM or **6** (0.06 mmol) in 0.5 mL MeOH was added the aldehyde (0.08 mmol) followed by 2 drops of AcOH. The resulting mixture was stirred for 30 min before NaBH_3_CN (0.1 mmol) was added. After stirring for 2 h at room temperature, the reaction was quenched with saturated NaHCO_3_ and extracted with DCM (3X). The combined organic layers were dried over Na_2_SO_4_, filtered, and concentrated. The residue was purified with silica gel column to provide desired product.

### General procedure B (Deprotection of bisubstrate inhibitors)

The protected inhibitor was dissolved in DCM (0.5 mL). The resulting solution was cooled to 0 °C and treated with TFA (0.5 mL). The reaction mixture was warmed to rt and stirred for 6 h. The solution was concentrated under reduced pressure, dissolved in water and purified by reverse HPLC to get the corresponding bisubstrate compound.

### ((3a*R*,4*R*,6*R*,6a*R*)-6-(6-chloro-9*H*-purin-9-yl)-2,2-dimethyltetrahydrofuro[3,4-*d*][1,3]dioxol-4-yl)methanol (2)

To a suspension of **1** (1.0 g, 3.4 mmol) in dry acetone (200 mL) was added p-TsOH monohydrate (6.6 g, 34.0 mmol) in one portion. The mixture was stirred at room temperature for 3 h. After completion of the reaction, ice-cold saturated NaHCO_3_ solution was added to the above mixture with stirring over 5 minutes. The volatiles were removed under reduced pressure and the crude was extracted with ethyl acetate (3X). The organic layers were combined, dried with sodium sulfate and concentrated to afford the crude product which was purified by column chromatography (dichloromethane: methanol 20:1) to afford **2** (1.1g, quant. yield) as a white solid.

### 2,2,2-trifluoro-*N*-(3-((9-((3a*R*,4*R*,6*R*,6a*R*)-6-(hydroxymethyl)-2,2-dimethyltetrahydrofuro[3,4-*d*][1,3]dioxol-4-yl)-9*H*-purin-6-yl)amino)propyl)acetamide (3)

To a solution of propane-1,3-diamine (450 mg, 6.1 mmol) and TEA (0.85 mL, 6.1 mmol) in EtOH (2 mL) was added slowly a solution of **2** (500 mg, 1.5 mmol) in EtOH (20 mL) and the reaction mixture was stirred at 60°C for 2 h. The solvent and excess of reagents were removed under reduced pressure to give the crude nucleoside. The crude nucleoside was dissolved in dry MeOH (20 mL) and TEA (0.42 mL, 3.1 mmol) and trifluoroacetic acid ethyl ester (1.1 mL, 9.1 mmol). The reaction mixture was stirred at room temperature for 12 h. The solvent was removed under reduced pressure and the crude product was purified by column chromatography (ethyl acetate 100 %) to give the protected nucleoside **3** (625 mg, 89 %) as a white solid. ^1^H NMR (500 MHz, CDCl_3_) δ 8.86 (s, 1H), 8.28 (s, 1H), 7.82 (s, 1H), 6.43 (t, *J* = 11.1 Hz, 2H), 5.85 (d, *J* = 4.9 Hz, 1H), 5.26 – 5.15 (m, 1H), 5.15 – 5.06 (m, 1H), 4.53 (s, 1H), 4.00 – 3.91 (m, 1H), 3.83 – 3.71 (m, 2H), 3.48 – 3.31 (m, 2H), 2.25 (s, 1H), 1.63 (s, 3H), 1.36 (s, 3H). ^13^C NMR (126 MHz, CDCl_3_) δ 157.48, 157.19, 155.80, 152.44, 147.39, 140.01, 117.75, 115.59, 114.12, 94.33, 86.12, 83.03, 81.67, 63.37, 37.67, 35.56, 29.41, 27.64, 25.23. LCMS (ESI) *m/z*: [M + H]^+^ 461.2

### *N*-(3-((9-((3a*R*,4*R*,6*R*,6a*R*)-6-((1,3-dioxoisoindolin-2-yl)methyl)-2,2-dimethyltetrahydrofuro[3,4-*d*][1,3]dioxol-4-yl)-9*H*-purin-6-yl)amino)propyl)-2,2,2-trifluoroacetamide (4)

To a solution of **3** (650 mg, 1.4 mmol) in dry THF (10 mL) was added phthalimide (249 mg, 1.7 mmol), triphenyl phosphine (740 mg, 2.8 mmol) and DIAD (566 mg, 2.8 mmol) at room temperature. The reaction mixture was stirred for 2 h. 10 mL of MeOH was added and stirring was continued for 15 min after which the crude was directly absorbed on silica gel and purified by column chromatography (ethyl acetate 100 %) to afford **4** (734 mg, 90 %) as a white solid. ^1^H NMR (500 MHz, CDCl_3_) δ 9.23 (s, 1H), 8.00 (s, 1H), 7.85 (s, 1H), 7.79 – 7.73 (m,2H), 7.67 (dd, *J* = 5.5, 3.0 Hz, 2H), 6.36 (s, 1H), 6.08 – 5.96 (m, 1H), 5.50 (d, *J* = 6.3 Hz, 1H), 5.23 (dd, *J* = 6.4, 3.5 Hz, 1H), 4.54 (q, *J* = 5.3 Hz, 1H), 4.06 – 3.91 (m, 2H), 3.72 (s, 1H), 3.39 (s, 2H), 1.85 (s, 2H), 1.56 (s, 3H), 1.36 (s, 3H). ^13^C NMR (126 MHz, CDCl_3_) δ 168.23, 157.48, 155.47, 152.58, 140.27, 134.12, 132.01, 123.39, 117.49, 115.20, 114.66, 90.86, 85.14, 84.18, 82.50, 39.54,37.01, 35.50, 29.73, 27.19, 25.48. LCMS (ESI) *m/z*: [M + H]^+^ 590.3

### *N*-(3-((9-((3a*R*,4*R*,6*R*,6a*R*)-6-(aminomethyl)-2,2-dimethyltetrahydrofuro[3,4-*d*][1,3]dioxol-4-yl)-9*H*-purin-6-yl)amino)propyl)-2,2,2-trifluoroacetamide (5)

To a solution of **4** (200 mg, 0.34 mmol) in ethanol was added hydrazine (34 mg, 0.67 mmol). The reaction was stirred at room temperature overnight. The reaction mixture was absorbed on silica gel and purified by column chromatography (DCM: MeOH 5:1) to afford **5** (98 mg, 63 %) as a white solid. ^1^H NMR (500 MHz, MeOD) δ 8.26 (s, 1H), 8.22 – 8.19 (m, 1H), 7.83 (dd, *J* = 7.9, 3.3 Hz, 1H), 6.21 – 6.10 (m, 1H), 5.47 – 5.38 (m, 1H), 5.13 – 5.03 (m, 1H), 4.40 – 4.26 (m, 1H), 3.67 (d, *J* = 38.2 Hz, 2H), 3.43 – 3.36 (m, 1H), 3.19 – 3.04 (m, 2H), 3.04 – 2.95 (m, 1H), 2.05 – 1.96 (m, 1H), 1.96 – 1.84 (m, 2H), 1.59 (s, 3H), 1.37 (s, 3H). ^13^C NMR (126 MHz, MeOD) δ 159.15, 153.18, 141.70, 133.49, 129.53, 126.72, 115.87, 91.54, 86.36, 85.17, 83.25, 43.50, 38.17, 27.45, 25.50. LCMS (ESI) *m/z*: [M + H]^+^ 460.3

### *tert*-butyl ((*S*)-1-((((3a*R*,4*R*,6*R*,6a*R*)-2,2-dimethyl-6-(6-((3-(2,2,2-trifluoroacetamido)propyl)amino)-9*H*-purin-9-yl)tetrahydrofuro[3,4-*d*][1,3]dioxol-4-yl)methyl)amino)-5,5-dimethyl-4-oxohexan-3-yl)carbamate (6)

Was prepared according to general procedure for reductive amination with **5** (200 mg, 0.4 mmol) and *tert*-butyl (*S*)-2-((*tert*-butoxycarbonyl)amino)-4-oxobutanoate (143 mg, 0.45 mmol) to afford **6** (209 mg, 67 %) as a white solid. ^1^H NMR (500 MHz, MeOD) δ 8.27 (s, 1H), 8.21 (s, 1H), 8.19 – 8.15 (m, 1H), 6.16 (d, *J* = 2.9 Hz, 1H), 5.44 (dd, *J* = 6.4, 3.0 Hz, 1H), 5.09 (dd, *J* = 6.5, 3.4 Hz, 1H), 4.42 – 4.34 (m, 1H), 4.06 (dd, *J* = 8.9, 4.8 Hz, 1H), 3.69 – 3.56 (m, 2H), 3.40 (t, *J* = 6.8 Hz, 2H), 3.10 (d, *J* = 5.8 Hz, 2H), 2.84 – 2.71 (m, 2H), 1.98 – 1.89 (m, 3H), 1.85 – 1.74 (m, 1H), 1.59 (s, 3H), 1.43 (s, 9H), 1.39 (s, 9H), 1.37 (s, 3H). ^13^C NMR (126 MHz, MeOD) δ 172.85, 159.09 - 158.51(m), 158.02, 156.30, 153.97, 141.56, 133.65, 126.62, 115.79, 91.85, 85.60, 84.88, 83.64, 82.90, 80.62, 53.87, 51.79, 46.90, 38.18, 31.37, 28.67, 28.22, 27.50, 25.60. LCMS (ESI) *m/z*: [M + H]^+^ 717.4

### *tert*-butyl ((*S*)-1-((3-(3-cyanophenyl)prop-2-yn-1-yl)(((3a*R*,4*R*,6*R*,6a*R*)-2,2-dimethyl-6-(6-((3-(2,2,2-trifluoroacetamido)propyl)amino)-9*H*-purin-9-yl)tetrahydrofuro[3,4-*d*][1,3]dioxol-4-yl)methyl)amino)-5,5-dimethyl-4-oxohexan-3-yl)carbamate (7)

Was prepared according to general procedure for reductive amination with **6** (170 mg, 0.24 mmol) and 3-(3-oxoprop-1-yn-1-yl)benzonitrile (44 mg, 0.28 mmol) to afford 6 (160 mg, 79 %) as a white solid. ^1^H NMR (500 MHz, MeOD) δ 8.24 (d, *J* = 2.7 Hz, 2H), 7.71 – 7.63 (m, 2H), 7.59 (d, *J* = 7.8 Hz, 1H), 7.48 (dd, *J* = 9.4, 6.2 Hz, 1H), 6.18 (d, *J* = 2.4 Hz, 1H), 5.49 (dd, *J* = 6.5, 2.5 Hz, 1H), 5.10 (dd, *J* = 6.4, 3.4 Hz, 1H), 4.43 – 4.35 (m, 1H), 4.15 – 4.02 (m, 1H), 3.69 – 3.58 (m, 3H), 3.39 (t, *J* = 6.9 Hz, 2H), 3.36 – 3.34 (m, 1H), 2.92 (dd, *J* = 13.5, 5.4 Hz, 1H), 2.84 (dd, *J* = 13.7, 7.5 Hz, 1H), 2.66 (t, *J* = 6.8 Hz, 2H), 1.98 – 1.87 (m, 3H), 1.59 (s, 3H), 1.43 (d, *J* = 2.4 Hz, 9H), 1.41 (s, 9H), 1.39 (s, 4H). ^13^C NMR (126 MHz, MeOD) δ 173.52, 159.11 −158.82 (m), 157.99, 156.20, 153.92, 141.53, 137.00, 135.89, 133.86, 132.60, 130.71, 126.56, 125.80, 118.97, 115.49, 113.79, 91.55, 87.90, 87.28, 85.04, 84.60, 84.17, 82.61, 80.45, 56.71, 54.16, 52.14, 44.33, 38.19, 30.27, 28.74, 28.28, 27.49, 25.65. LCMS (ESI) *m/z*: [M + H]^+^ 856.5

### *tert*-butyl ((*S*)-1-((3-(3-carbamoylphenyl)prop-2-yn-1-yl)(((3a*R*,4*R*,6*R*,6a*R*)-2,2-dimethyl-6-(6-((3-(2,2,2-trifluoroacetamido)propyl)amino)-9*H*-purin-9-yl)tetrahydrofuro[3,4-*d*][1,3]dioxol-4-yl)methyl)amino)-5,5-dimethyl-4-oxohexan-3-yl)carbamate (8)

A solution of **7** (160 mg, 0.18 mmol) and K_2_CO_3_ (103 mg, 0.75 mmol) in DMSO (3 mL) was cooled to 0 °C and then treated with hydrogen peroxide (38 mg, 1.12 mmol). The reaction mixture was warmed to room temperature and stirred for 3 h. The reaction mixture was diluted with water and extracted with ethyl acetate (3X). The organic layers were combined, dried with sodium sulfate and concentrated to afford the crude product which was purified by column chromatography (DCM: MeOH 10:1) to afford **8** (163 mg, quant. yield) as a white solid. ^1^H NMR (500 MHz, CDCl_3_) δ 9.08 (s, 1H), 8.30 (s, 1H), 7.99 (s, 1H), 7.88 – 7.73 (m, 2H), 7.46 (d, *J* = 7.7 Hz, 1H), 7.35 (t, *J* = 7.8 Hz, 1H), 6.85 (s, 1H), 6.40 (t, *J* = 58.7 Hz, 2H), 6.10 (s, 1H), 5.59 (t, *J* = 17.6 Hz, 1H), 5.47 (d, *J* = 6.3 Hz, 1H), 5.07 – 4.82 (m, 1H), 4.50 – 4.36 (m, 1H), 3.81 – 3.54 (m, 4H), 3.44 – 3.27 (m, 2H), 2.82 (p, *J* = 8.9, 7.2 Hz, 2H), 2.64 (dd, *J* = 16.3, 9.0 Hz, 2H), 2.02 – 1.74 (m, 4H), 1.61 (s, 3H), 1.42 (s, 9H), 1.40 (s, 9H), 1.39 (s, 3H). ^13^C NMR (126 MHz, CDCl_3_) δ 171.85, 169.12, 157.54 −157.25 (m), 155.68, 155.53, 153.02, 139.79, 134.89, 133.70, 130.80, 128.74, 127.37, 123.48, 117.48, 115.19, 114.62, 90.97, 86.38, 85.25, 84.64, 84.07, 83.37, 81.99, 79.74, 55.84, 52.94, 50.87, 43.71, 30.06, 29.42, 28.44, 28.08, 27.24, 25.52. LCMS (ESI) *m/z*: [M + H]^+^ 874.4

### *tert*-butyl ((*S*)-1-((((3a*R*,4*R*,6*R*,6a*R*)-6-(6-((3-aminopropyl)amino)-9*H*-purin-9-yl)-2,2-dimethyltetrahydrofuro[3,4-*d*][1,3]dioxol-4-yl)methyl)(3-(3-carbamoylphenyl)prop-2-yn-1-yl)amino)-5,5-dimethyl-4-oxohexan-3-yl)carbamate (9)

Nucleoside **8** (160 mg, 0.18 mmol) was dissolved in a mixture of MeOH (6 mL) and aqueous NH_3_ solution (6 mL, 33%) and stirred at room temperature for 12 h. The solvents were removed *in vacuo*, the crude product was dissolved in DCM (10 mL) and the organic layer washed with aqueous TEA solution. Removal of the solvent yielded the protected AdoHcy analog **9** (123 mg, 77%) as a clear oil. ^1^H NMR (500 MHz, CDCl_3_) δ 8.32 (s, 1H), 7.93 (s, 1H), 7.82 (d, *J* = 9.7 Hz, 2H), 7.47 – 7.42 (m, 1H), 7.34 (t, *J* = 7.7 Hz, 1H), 6.08 (d, *J* = 1.9 Hz, 1H), 5.59 (d, *J* = 8.1 Hz, 1H), 5.46 (d, *J* = 6.5 Hz, 1H), 5.03 – 4.94 (m, 1H), 4.44 – 4.35 (m, 1H), 4.29 – 4.17 (m, 1H), 3.78 – 3.52 (m, 4H), 2.88 – 2.77 (m, 4H), 2.67 – 2.56 (m, 2H), 1.86 – 1.73 (m, 4H), 1.59 (s, 4H), 1.42 (s, 9H), 1.40 (s, 9H), 1.38 (s, 3H). ^13^C NMR (126 MHz, CDCl_3_) δ 171.84, 169.02, 155.64, 155.06, 153.37, 139.28, 134.79, 133.77, 130.68, 128.73, 127.68, 123.32, 120.42, 114.45, 90.93, 86.71, 85.25, 84.62, 84.12, 83.47, 81.93, 79.68, 55.82, 52.92, 50.93, 43.86, 39.82, 30.01, 28.45, 28.10, 27.23, 25.54. LCMS (ESI) *m/z*: [M + H]^+^ 778.5

### (*S*)-2-amino-4-((((2*R*,3*S*,4*R*,5*R*)-5-(6-((3-aminopropyl)amino)-9*H*-purin-9-yl)-3,4-dihydroxytetrahydrofuran-2-yl)methyl)(3-(3-carbamoylphenyl)prop-2-yn-1-yl)amino)butanoic acid (10)

Was prepared according to general procedure for deprotection of bisubstrate inhibitor **9** (20 mg, 0.026 mmol) to afford **10** (6 mg, 37 %) as a white solid. ^1^H NMR (500 MHz, Deuterium Oxide) δ 8.37 (s, 1H), 8.23 (d, *J* = 3.5 Hz, 1H), 7.87 (d, *J* = 7.4 Hz, 1H), 7.59 (s, 1H), 7.52 – 7.41 (m, 2H), 6.19 (d, *J* = 3.5 Hz, 1H), 4.51 (dd, *J* = 16.9, 3.4 Hz, 3H), 4.36 (dd, *J* = 17.0, 3.5 Hz, 2H), 4.06 – 3.98 (m, 2H), 3.94 – 3.89 (m, 1H), 3.85 – 3.79 (m, 1H), 3.63 (d, *J* = 7.2 Hz, 2H), 3.48 (s, 2H), 3.13 – 3.05 (m, 2H), 2.49 – 2.37 (m, 1H), 2.27 (d, *J* = 12.6 Hz, 1H), 2.10 – 1.99 (m, 2H). HRMS *m/z* calcd for C_27_H_36_N_9_O_6_ [M + H]^+^: 582.2783; found: 582.2783

### (*S*)-2-amino-4-((3-(3-carbamoylphenyl)prop-2-yn-1-yl)(((2*R*,3*S*,4*R*,5*R*)-5-(6-((3-(3’,6’-dihydroxy-3-oxo-3*H*-spiro[isobenzofuran-1,9’-xanthene]-5-carboxamido)propyl)amino)-9*H*-purin-9-yl)-3,4-dihydroxytetrahydrofuran-2-yl)methyl)amino)butanoic acid (11)

To a solution of **9** (30 mg, 0.038 mmol) in DMF (1 mL) was added DIC (10 mg, 0.077 mmol) and the reaction was stirred for 30 mins before the addition of HOBt (7.8 mg, 0.058 mmol), 5-Carboxyfluorescein (17 mg, 0.046 mmol) and DIPEA (10 mg, 0.077 mmol). The reaction was stirred at room temperature overnight. The reaction mixture was quenched with water and extracted with ethyl acetate (3X). The organic layers were combined, washed with brine, dried with sodium sulfate and concentrated to afford the crude product which was then subjected to the general procedure for deprotection of bisubstrate inhibitor to afford **11** (12 mg, 29 %) as a yellow solid. ^1^H NMR (500 MHz, DMSO-*d*_6_) δ 10.26 (s, 1H), 8.91 (s, 1H), 8.45 (s, 1H), 8.37 (s, 1H), 8.24 (s, 2H), 8.08 (s, 1H), 7.94 (s, 1H), 7.85 (d, *J* = 7.3 Hz, 1H), 7.53 (s, 1H), 7.44 (s, 1H), 7.37 (d, *J* = 7.6 Hz, 1H), 6.71 (s, 2H), 6.55 (d, *J* = 14.1 Hz, 4H), 5.91 (s, 1H), 5.54 (s, 1H), 4.66 (s, 1H), 4.15 (d, *J* = 22.8 Hz, 2H), 3.94 (s, 1H), 2.95 (s, 2H), 2.83 (s, 2H), 2.02 (s, 1H), 1.91 (d, *J* = 38.7 Hz, 3H), 1.20 (d, *J* = 17.7 Hz, 1H). ^13^C NMR (126 MHz, DMSO-*d*_6_) δ 170.82, 168.19, 167.02, 164.68, 159.61, 154.61, 152.61, 151.76, 139.69, 136.31, 134.64, 134.53, 134.04, 130.40, 129.07, 128.67, 127.51, 126.41, 124.23, 123.20, 122.25, 119.59, 112.68, 108.98, 102.27, 87.80, 84.87, 83.29, 81.88, 72.72, 71.82, 55.41, 51.09, 49.88, 42.74, 37.54, 37.06, 28.92, 27.14. HRMS *m/z* calcd for C_48_H_46_N_9_O_12_ [M + H]^+^: 940.3260; found: 940.3259

## 1. Abbreviations used

AdoHcy: S-adenosylhomocysteine
AdoMet: S-adenosylmethionine
SAHH: S-adenosylhomocysteine hydrolase
NAM: nicotinamide
MNAM: N1-methylnicotinamide
NNMT: nicotinamide N-methyltransferase
HPLC: High-Pressure Liquid chromatography
FP: fluorescence polarization
HTS: high-throughput screening
TFA: trifluoroacetic acid
NMR: Nuclear magnetic resonance
*K*_*i*_: inhibitory constant
*K*_*d*_: dissociation constant
DMSO: dimethyl sulfoxide.

## Acknowledgements

We appreciate Dr. Dongxing Chen for his assistance with the enzymatic activity assay and Krystal Diaz for her feedback. The authors acknowledge the support from NIH grants R01GM117275 (R.H.), U01CA214649 (R.H.), and P30 CA023168 (Purdue University Center for Cancer Research). We also thank supports from the Department of Medicinal Chemistry and Molecular Pharmacology (R.H.) at Purdue University.

## Note

The authors declare no competing financial interest.

## Supplementary data

Supplementary data to this article is attached.

