## supplemental file for "Development of fluorescence polarization-based competition assay for nicotinamide *N*-methyltransferase"

#### Table of Contents

A. NMR spectra of compounds **3-9** S2

B. NMR, HR-MS and HPLC spectra of compounds **10 and II138** S16

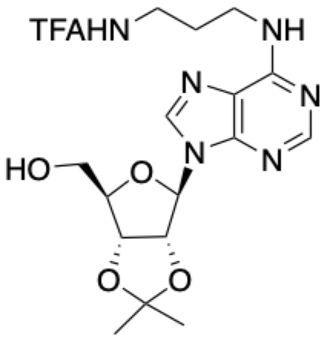
TFAHN
NH
N
N
N
N
HO
O
O
O

**3**

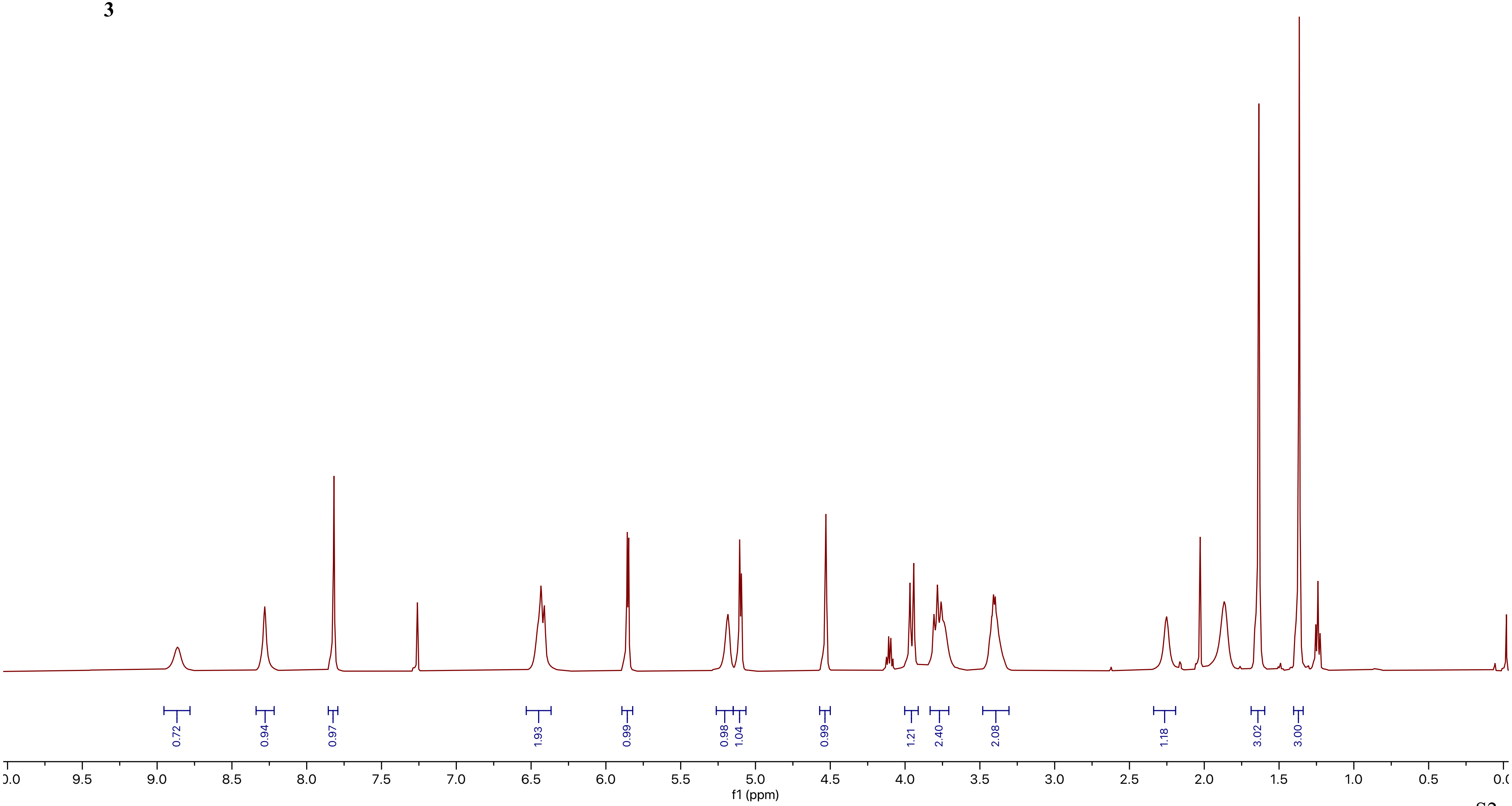
0.72
0.94
0.97
1.93
0.99
0.98
1.04
0.99
1.21
2.40
2.08
1.18
3.02
3.00
9.5
9.0
8.5
8.0
7.5
7.0
6.5
6.0
5.5
5.0
4.5
4.0
3.5
3.0
2.5
2.0
1.5
1.0
0.5
f1 (ppm)

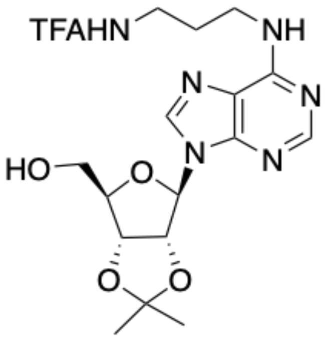

TFAHN
NH
N
N
N
N
HO
O
O
O


**3**

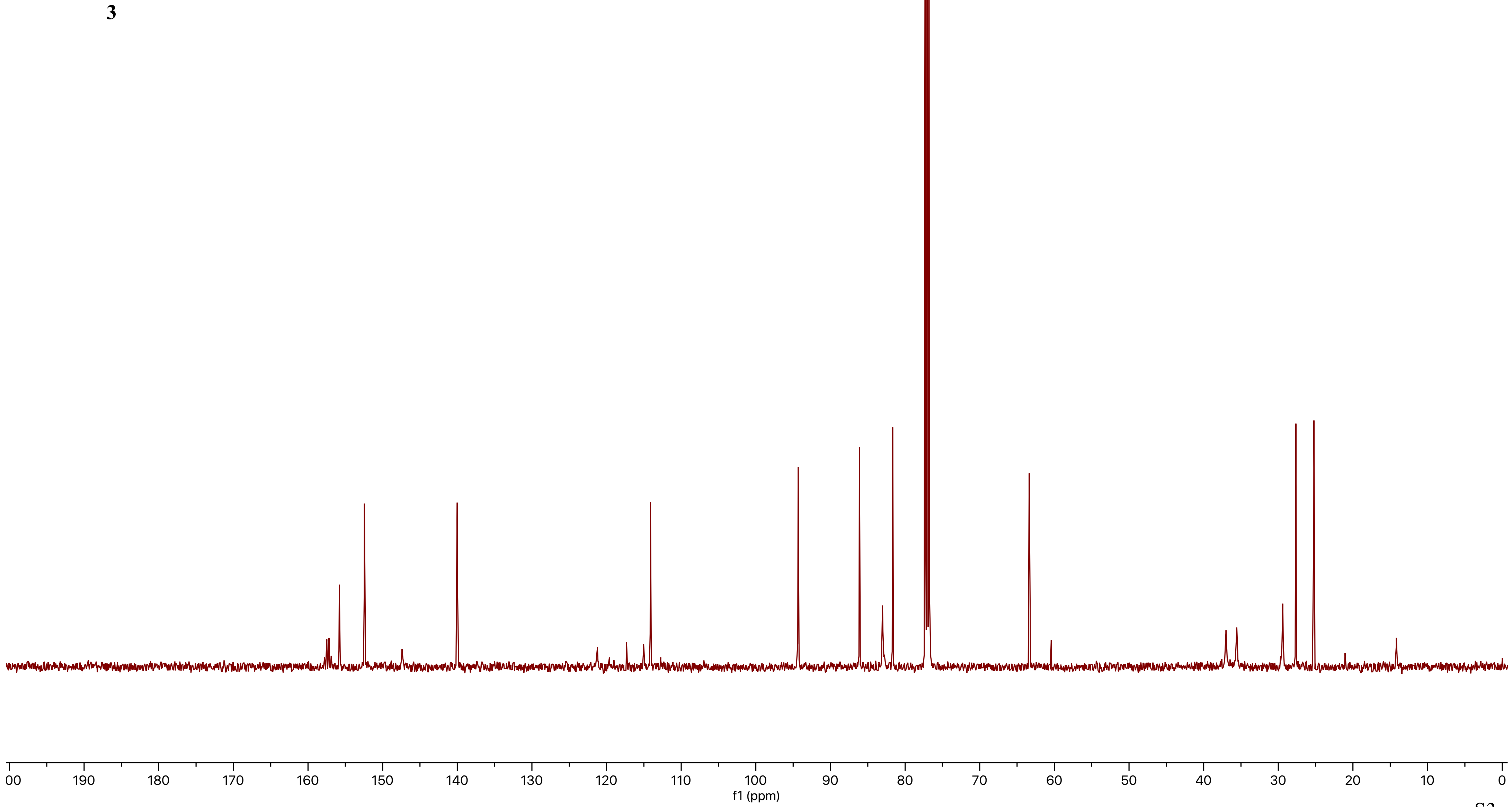

157.48
157.19
155.80
152.44
147.39
140.01
117.75
115.59
114.12
94.33
86.12
83.03
81.67
63.37
37.67
35.56
29.41
27.64
25.23
00 190 180 170 160 150 140 130 120 110 100 90 80 70 60 50 40 30 20 10 0
f1 (ppm)

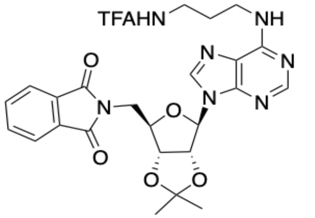
TFAHN
NH
N
N
N
N
N
O
O
O
O
O

**4**

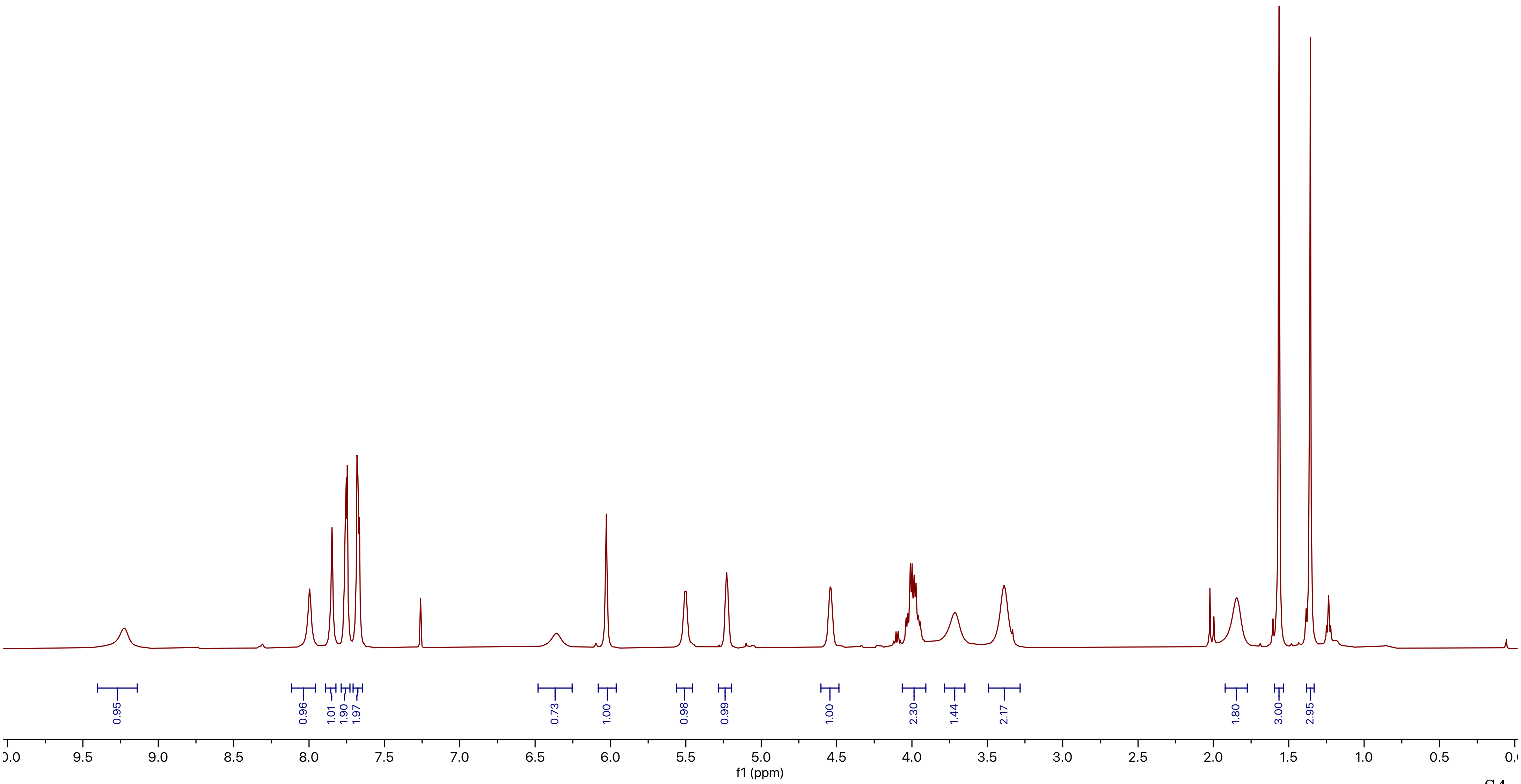
0.95
0.96
1.01
1.90
1.97
0.73
1.00
0.98
0.99
1.00
2.30
1.44
2.17
1.80
3.00
2.95
9.5
9.0
8.5
8.0
7.5
7.0
6.5
6.0
5.5
5.0
4.5
4.0
3.5
3.0
2.5
2.0
1.5
1.0
0.5
f1 (ppm)

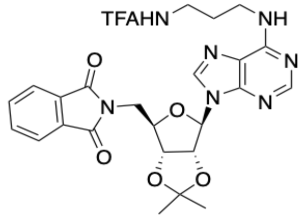
TFAHN
NH
N
N
N
N
O
O
N
O
O
O
4

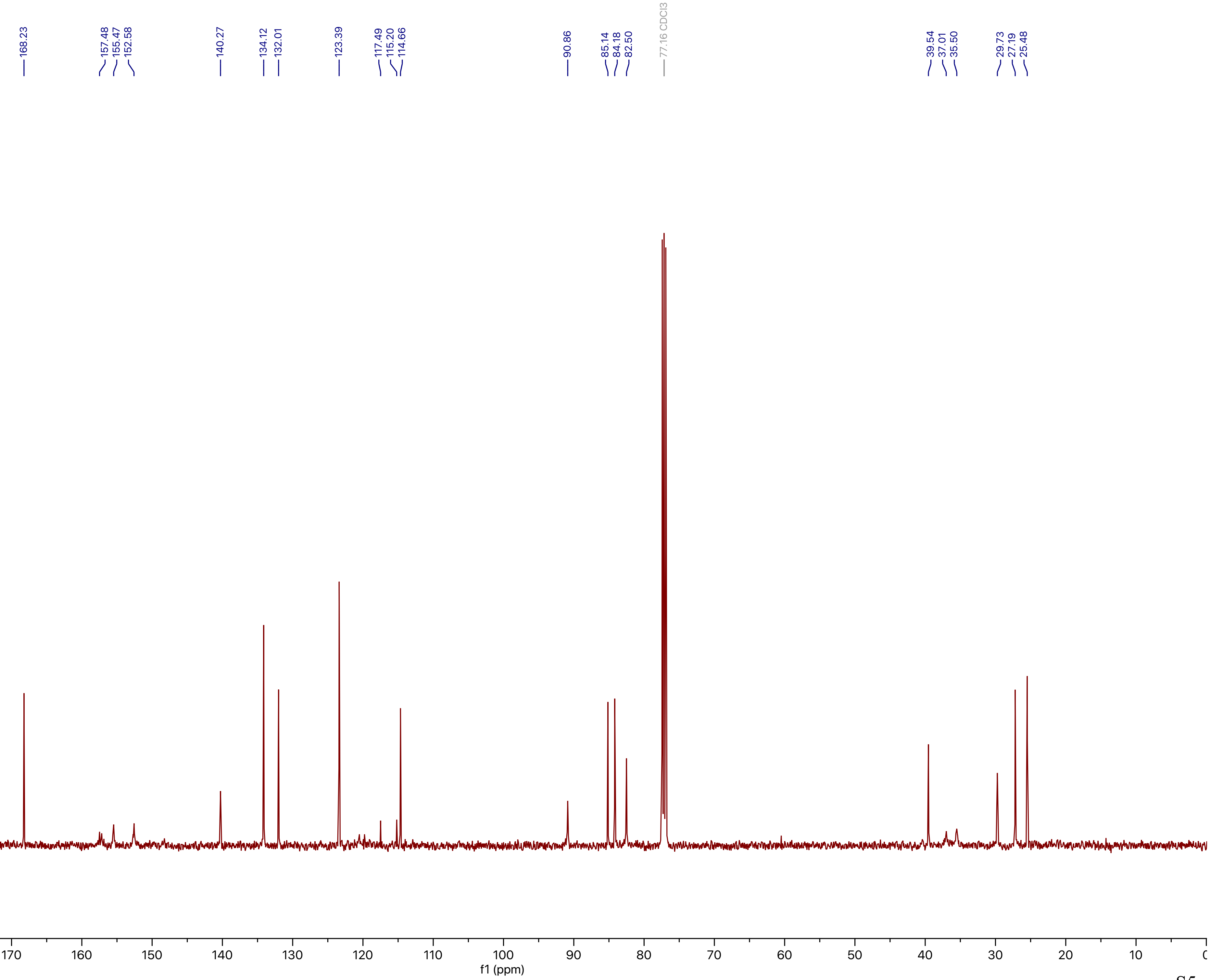
168.23
157.48
155.47
152.58
140.27
134.12
132.01
123.39
117.49
115.20
114.66
90.86
85.14
84.18
82.50
77.16 CDCl3
39.54
37.01
35.50
29.73
27.19
25.48
190
180
170
160
150
140
130
120
110
100
90
80
70
60
50
40
30
20
10
f1 (ppm)

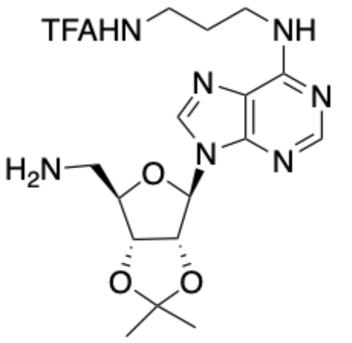
TFAHN
NH
N
N
N
N
H2N
O
O
O

**5**

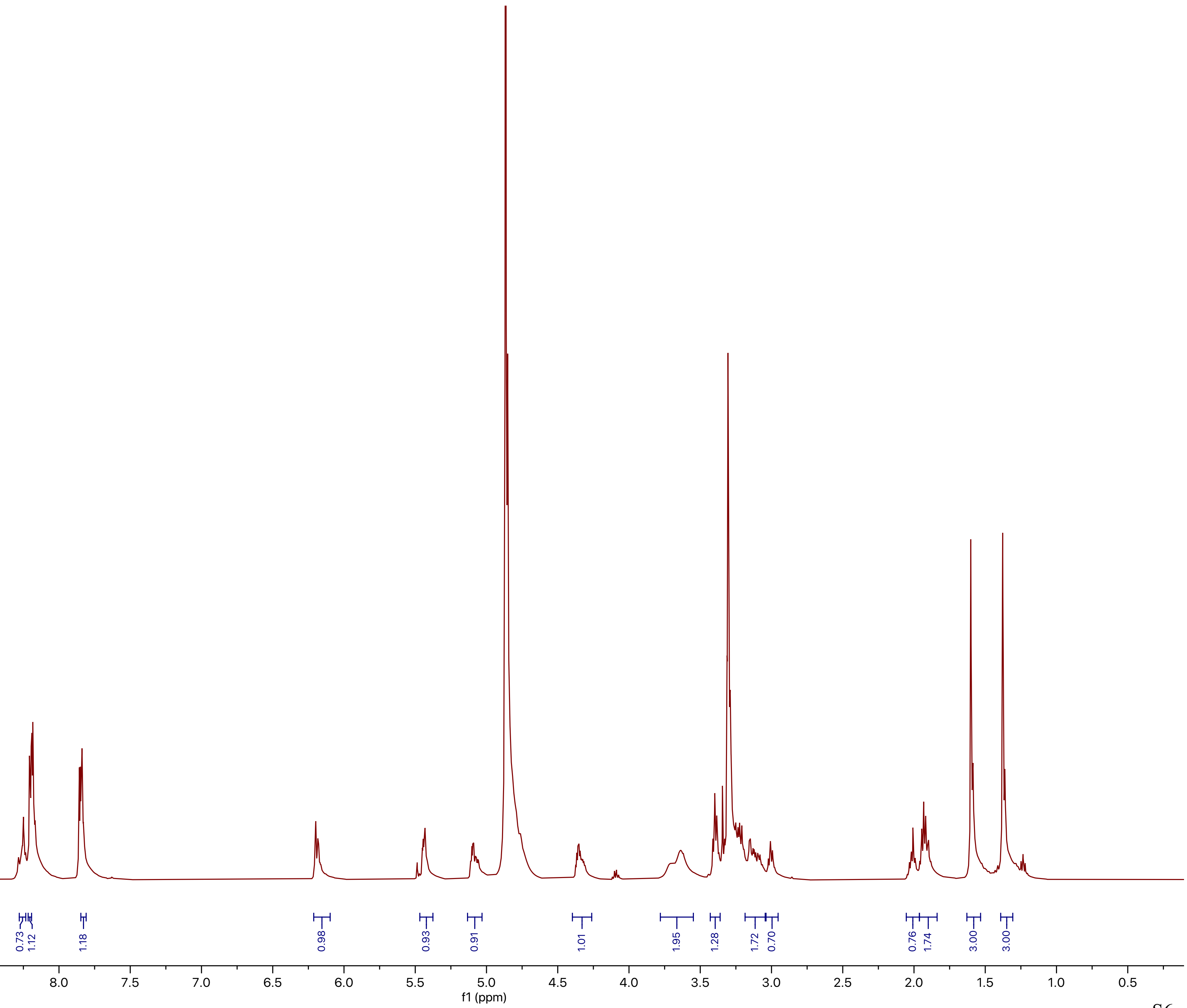
0.73
1.12
1.18
0.98
0.93
0.91
1.01
1.95
1.28
1.72
0.70
0.76
1.74
3.00
3.00
10.0 9.5 9.0 8.5 8.0 7.5 7.0 6.5 6.0 5.5 5.0 4.5 4.0 3.5 3.0 2.5 2.0 1.5 1.0 0.5
f1 (ppm)

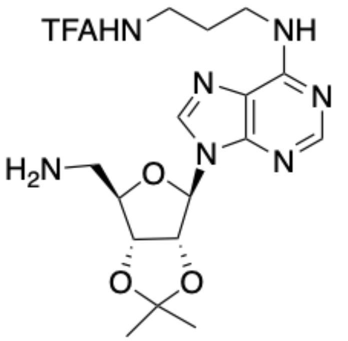
TFAHN
NH
N
N
N
N
H2N
O
O
O

**5**

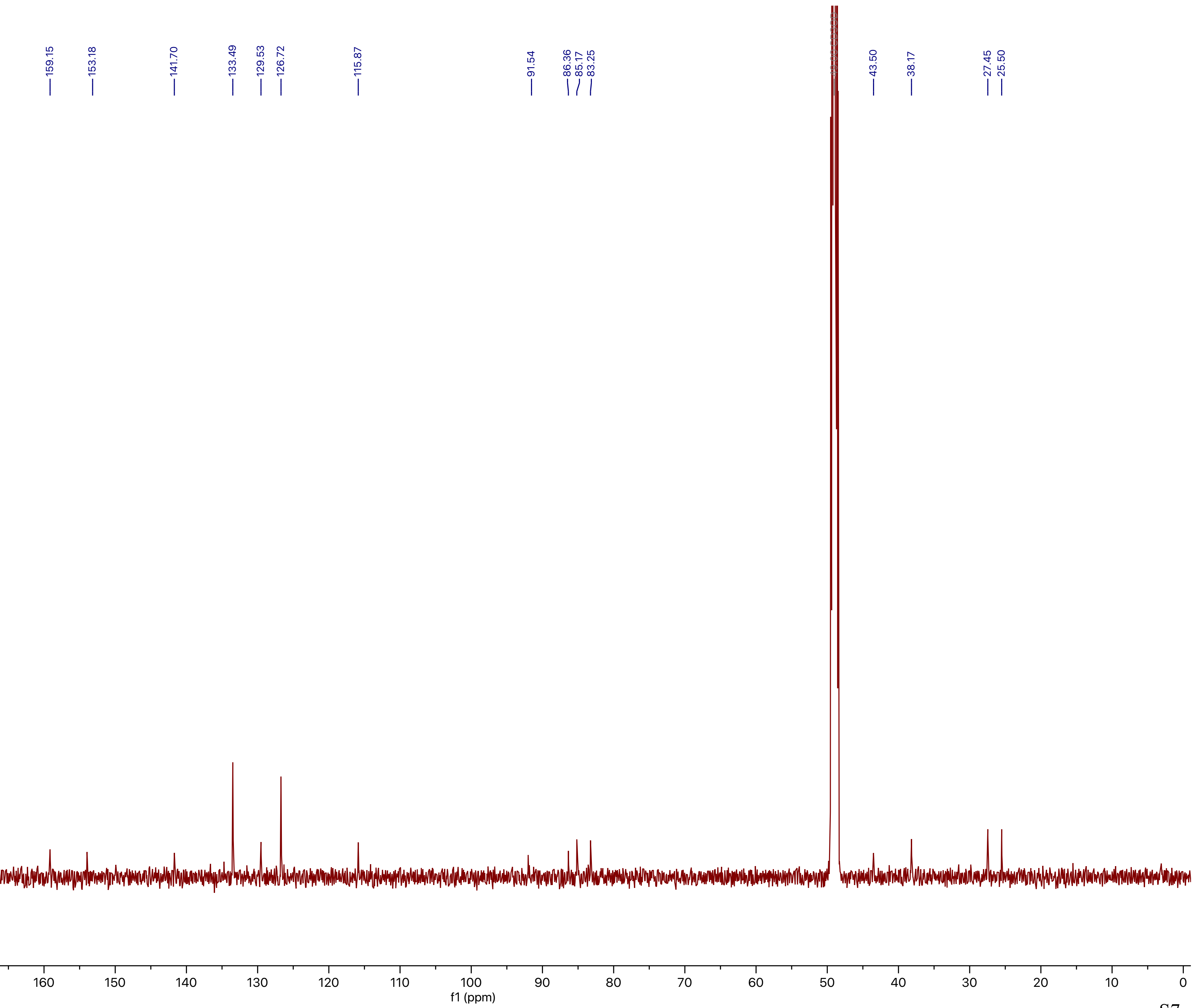
159.15
153.18
141.70
133.49
129.53
126.72
115.87
91.54
86.36
85.17
83.25
43.50
38.17
27.45
25.50
00
190
180
170
160
150
140
130
120
110
100
90
80
70
60
50
40
30
20
10
0
f1 (ppm)

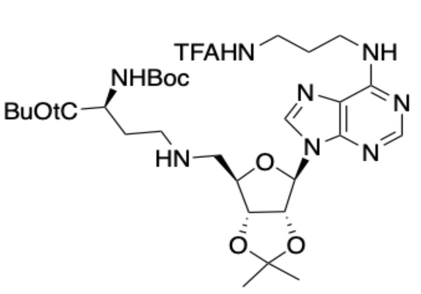
TFAHN
NH
NHBoc
BuOtC
HN

**6**

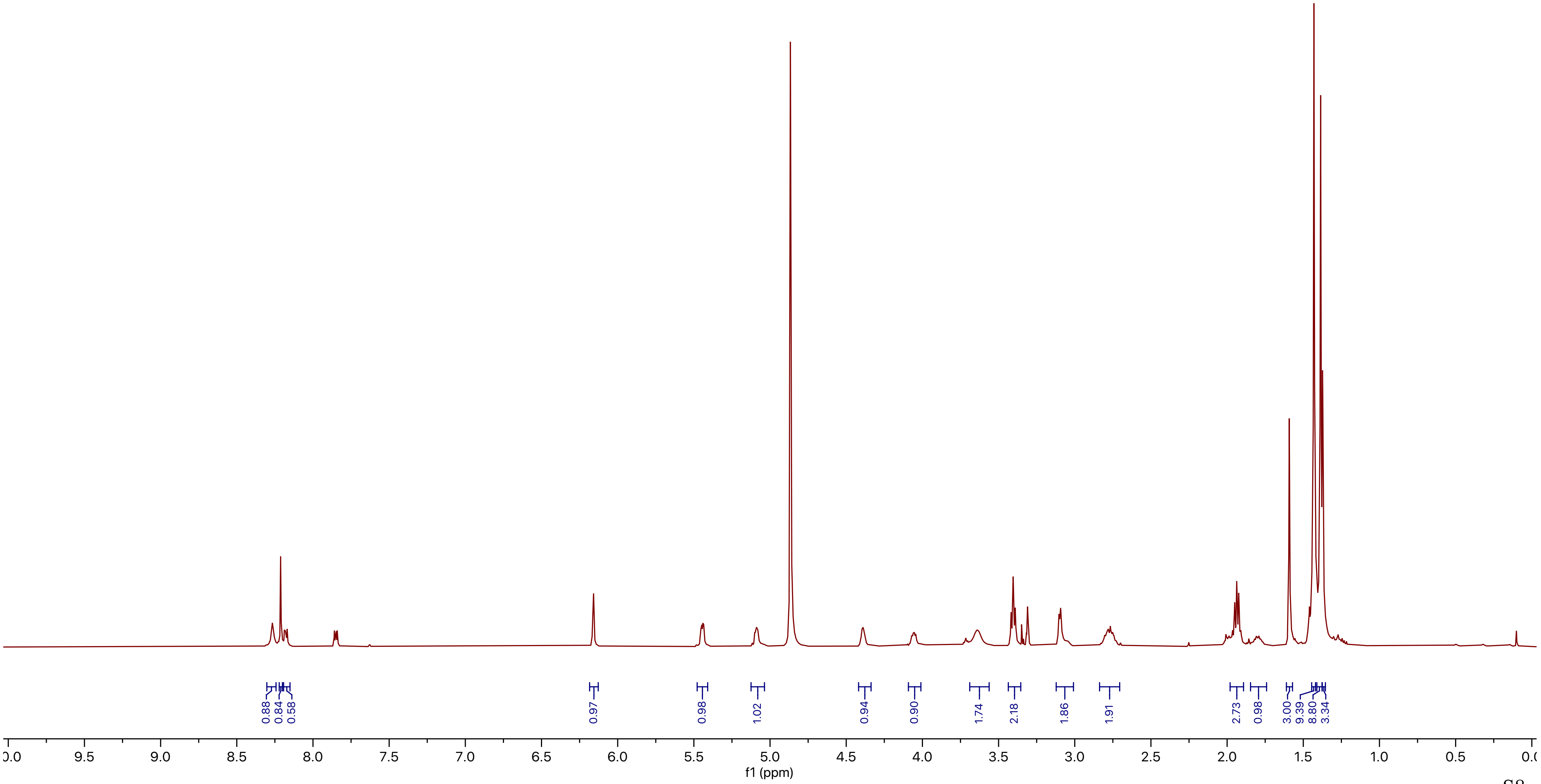
0.88
0.84
0.58
0.97
0.98
1.02
0.94
0.90
1.74
2.18
1.86
1.91
2.73
0.98
3.00
9.39
8.80
3.34
9.5
9.0
8.5
8.0
7.5
7.0
6.5
6.0
5.5
5.0
4.5
4.0
3.5
3.0
2.5
2.0
1.5
1.0
0.5
f1 (ppm)

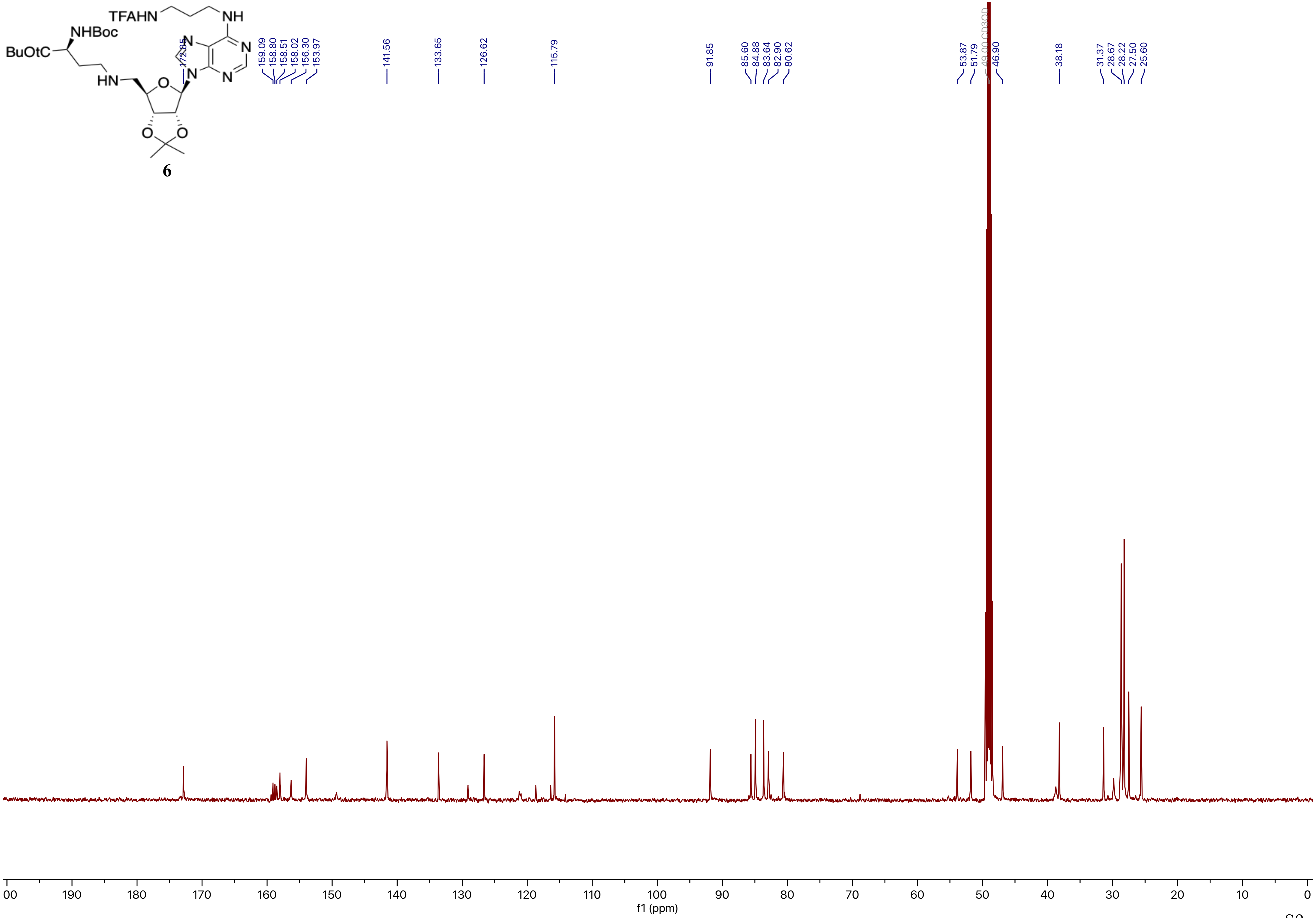
TFAHN
NH
NHBoc
BuOtC
HN
N
N
N
N
O
O
O
6
172.85
159.09
158.80
158.51
158.02
156.30
153.97
141.56
133.65
126.62
115.79
91.85
85.60
84.88
83.64
82.90
80.62
53.87
51.79
49.00 CD3OD
46.90
38.18
31.37
28.67
28.22
27.50
25.60
00
190
180
170
160
150
140
130
120
110
100
90
80
70
60
50
40
30
20
10
0
f1 (ppm)

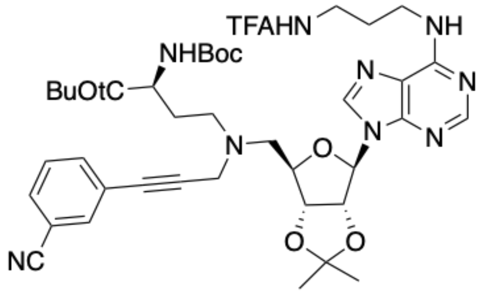
TFAHN
NH
NHBoc
BuOtC
N
N
N
N
N
O
O
O
NC

7

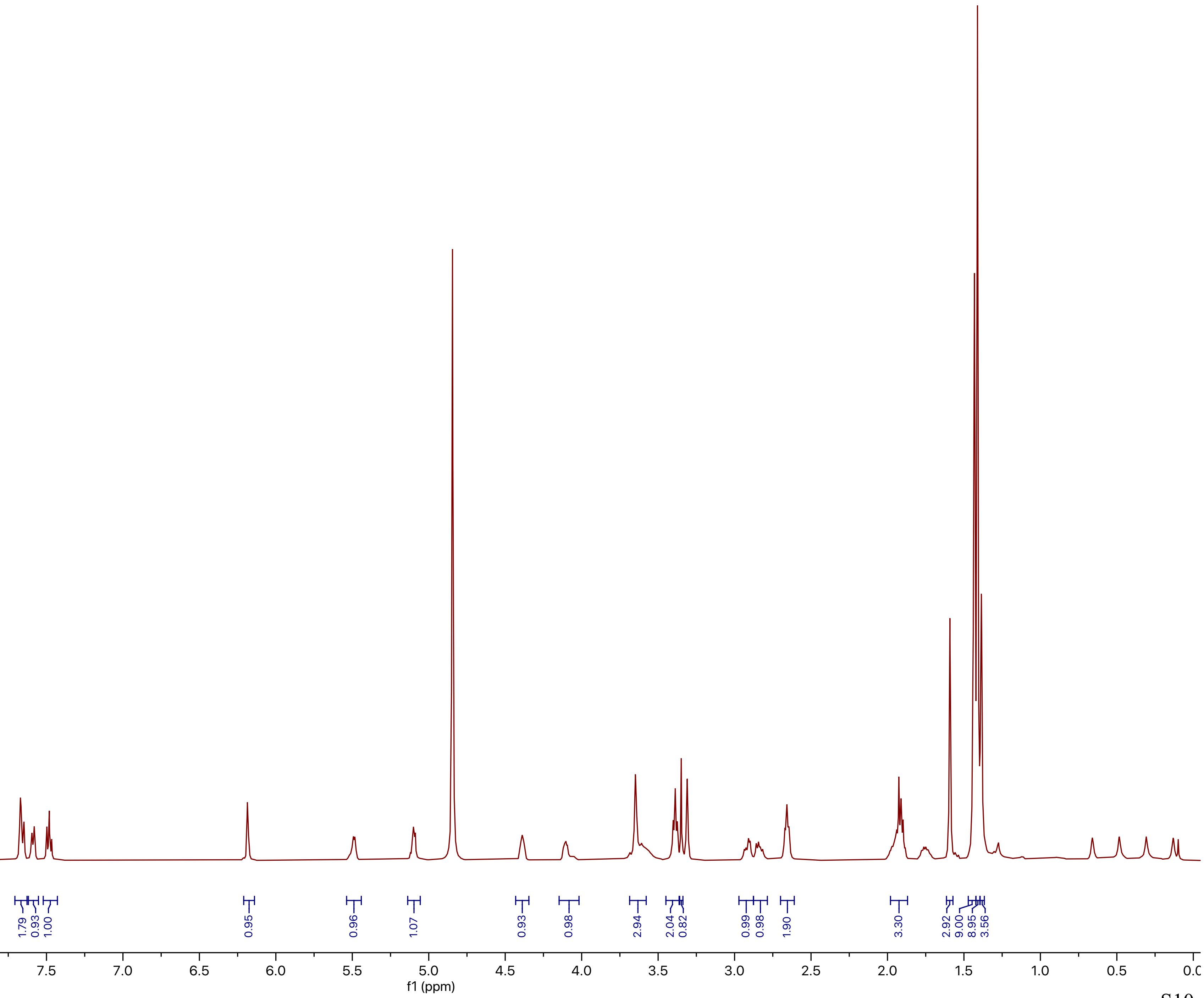
1.86
1.79
0.93
1.00
0.95
0.96
1.07
0.93
0.98
2.94
2.04
0.82
0.99
0.98
1.90
3.30
2.92
9.00
8.95
3.56
9.5
9.0
8.5
8.0
7.5
7.0
6.5
6.0
5.5
5.0
4.5
4.0
3.5
3.0
2.5
2.0
1.5
1.0
0.5
f1 (ppm)

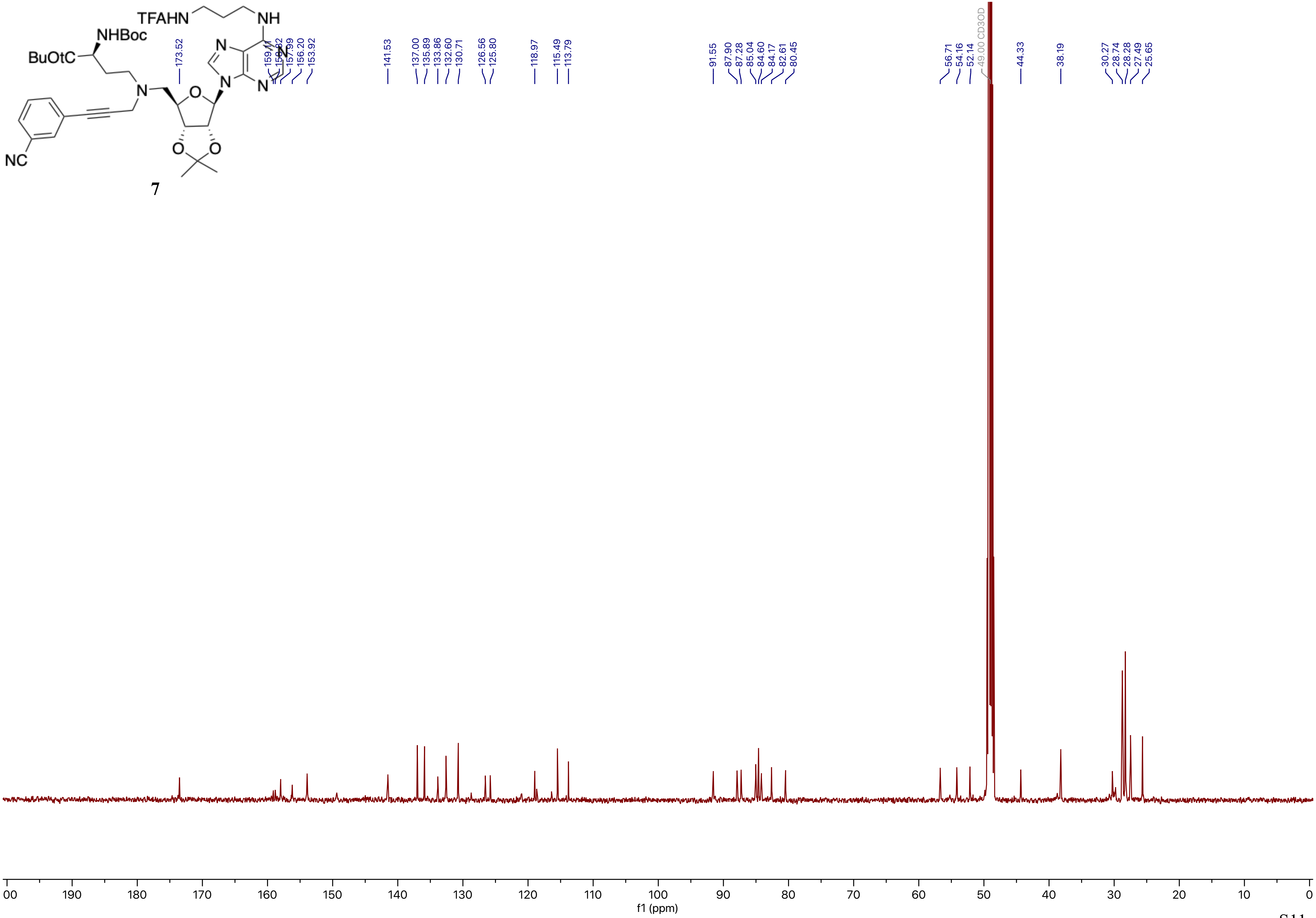
TFAHN
NH
NHBoc
BuOtC
NC
7
173.52
159.11
157.99
156.20
153.92
141.53
137.00
135.89
133.86
132.60
130.71
126.56
125.80
118.97
115.49
113.79
91.55
87.90
87.28
85.04
84.60
84.17
82.61
80.45
56.71
54.16
52.14
49.00 CD3OD
44.33
38.19
30.27
28.74
28.28
27.49
25.65
f1 (ppm)

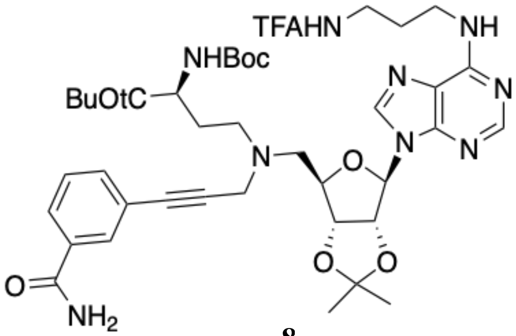
TFAHN
NH
NHBoc
BuOtC
N
N
N
N
N
O
O
O
O
NH2

**8**

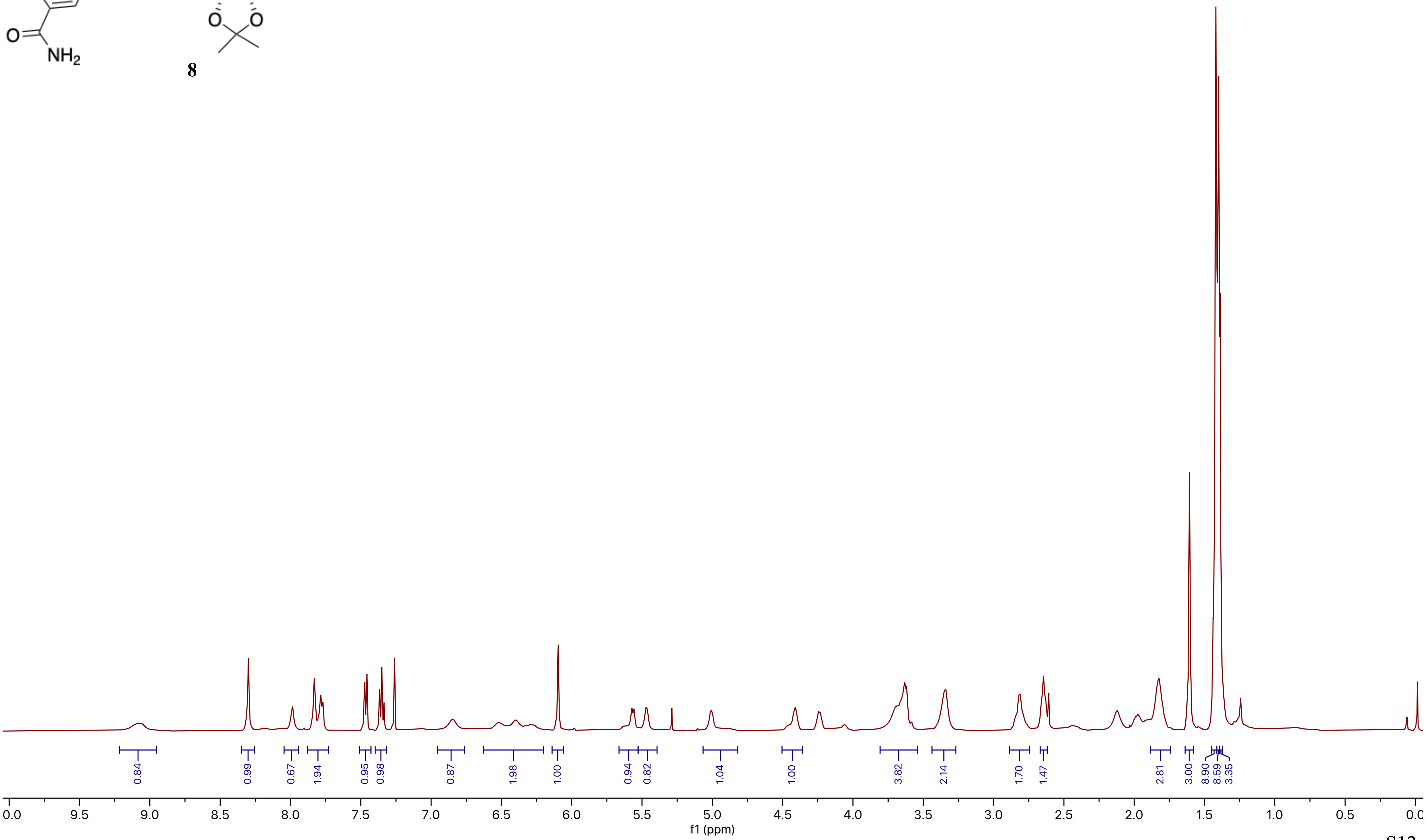
0.84
0.99
0.67
1.94
0.95
0.98
0.87
1.98
1.00
0.94
0.82
1.04
1.00
3.82
2.14
1.70
1.47
2.81
3.00
8.90
8.59
3.35
10.0
9.5
9.0
8.5
8.0
7.5
7.0
6.5
6.0
5.5
5.0
4.5
4.0
3.5
3.0
2.5
2.0
1.5
1.0
0.5
0.0
f1 (ppm)

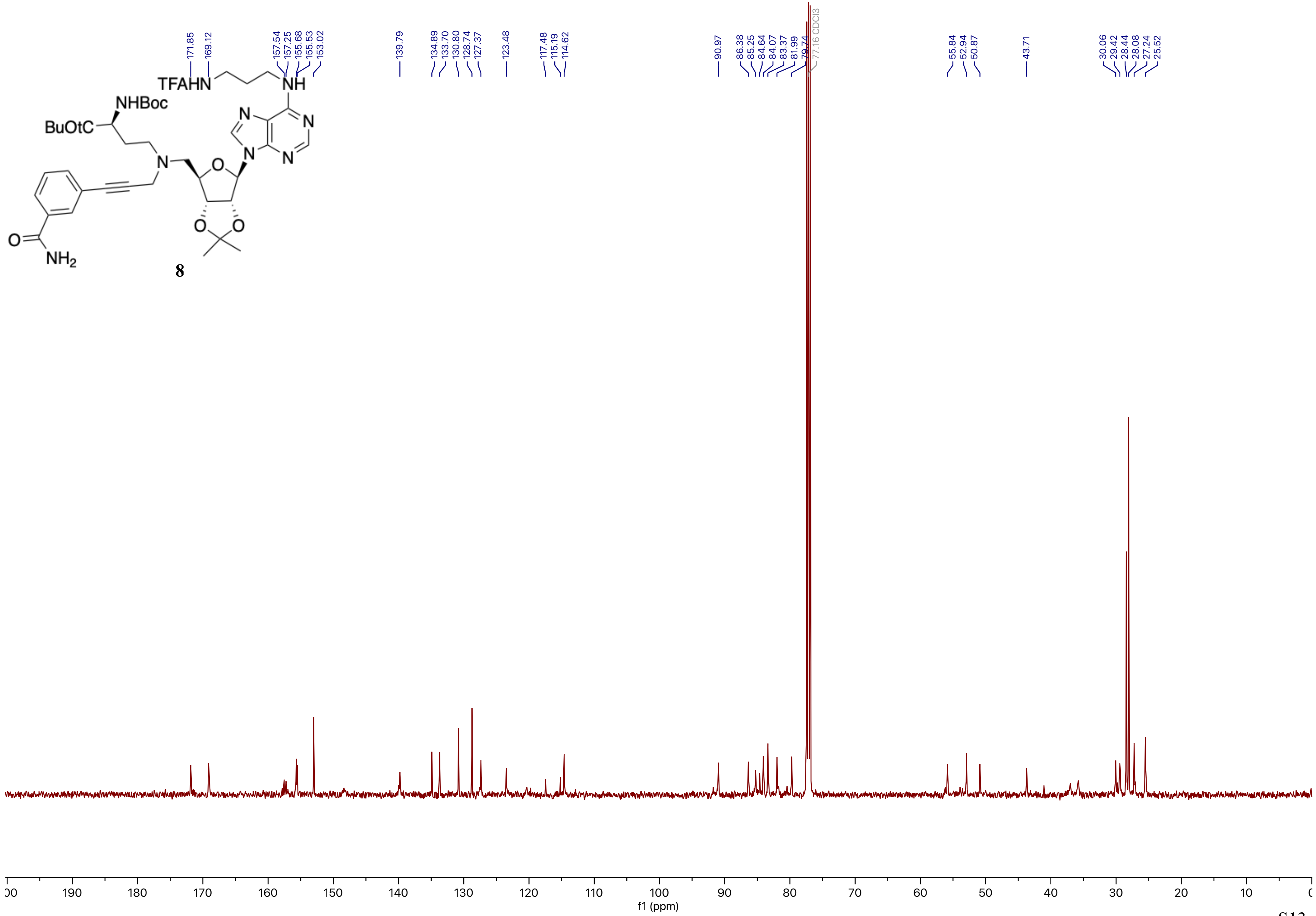
171.85
169.12
157.54
157.25
155.68
155.53
153.02
139.79
134.89
133.70
130.80
128.74
127.37
123.48
117.48
115.19
114.62
90.97
86.38
85.25
84.64
84.07
83.37
81.99
79.74
77.16 CDCl3
55.84
52.94
50.87
43.71
30.06
29.42
28.44
28.08
27.24
25.52
TFAHN
NH
NHBoc
BuOtC
N
O
NH2
8
f1 (ppm)

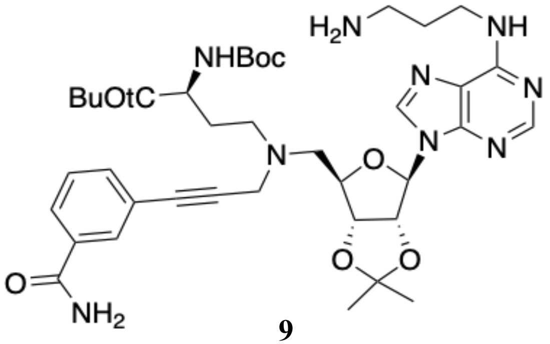
H2N
NH
NHBoc
BuOtC
N
O
NH2

**9**

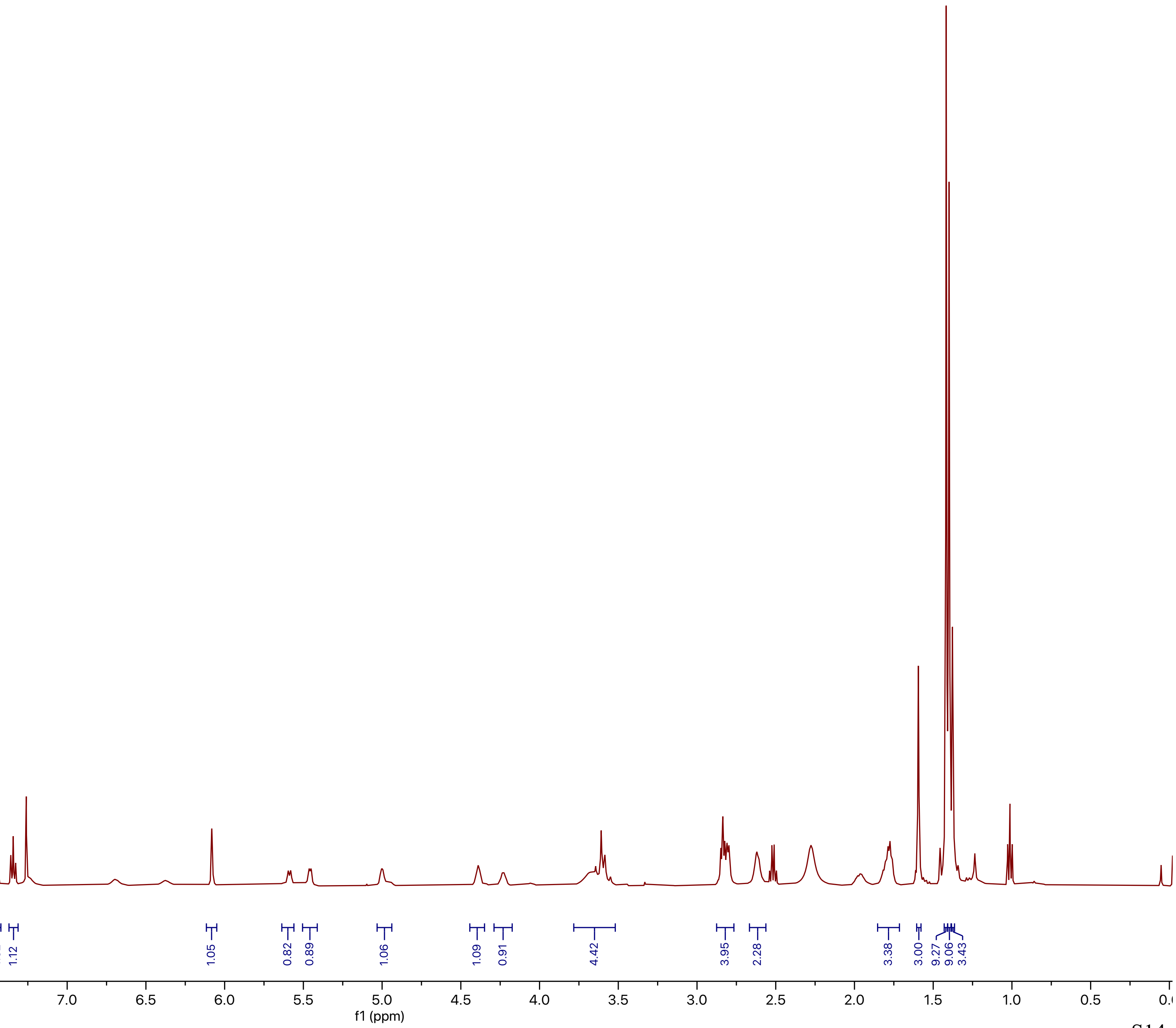
1.01
0.86
2.06
1.02
1.12
1.05
0.82
0.89
1.06
1.09
0.91
4.42
3.95
2.28
3.38
3.00
9.27
9.06
3.43
0.0
9.5
9.0
8.5
8.0
7.5
7.0
6.5
6.0
5.5
5.0
4.5
4.0
3.5
3.0
2.5
2.0
1.5
1.0
0.5
f1 (ppm)

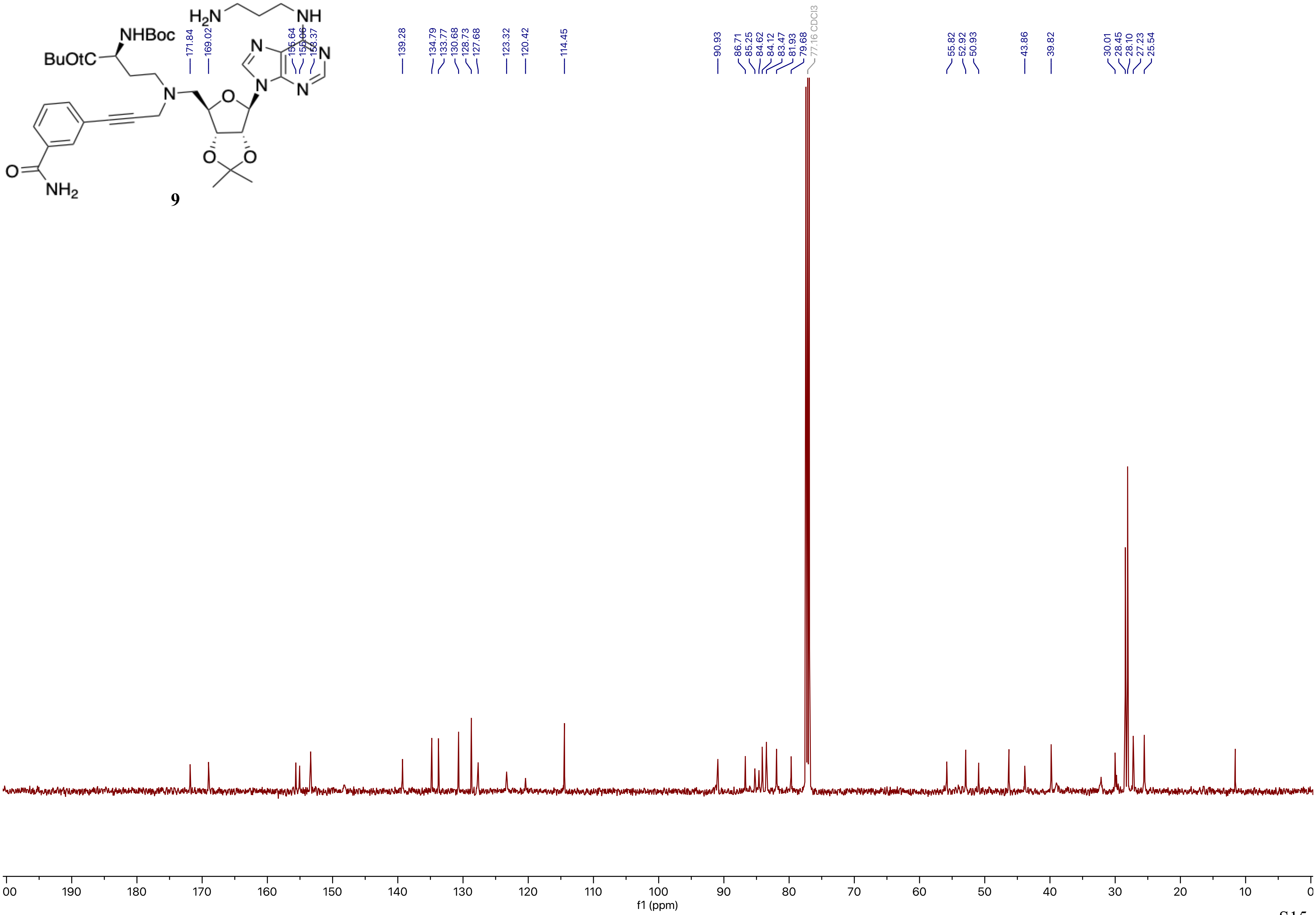

H2N
NH
NHBoc
BuOtC
N
O
NH2
171.84
169.02
155.64
153.37
139.28
134.79
133.77
130.68
128.73
127.68
123.32
120.42
114.45
90.93
86.71
85.25
84.62
84.12
83.47
81.93
79.68
77.16 CDCl3
55.82
52.92
50.93
43.86
39.82
30.01
28.45
28.10
27.23
25.54
00
190
180
170
160
150
140
130
120
110
100
90
80
70
60
50
40
30
20
10
0
f1 (ppm)


**9**

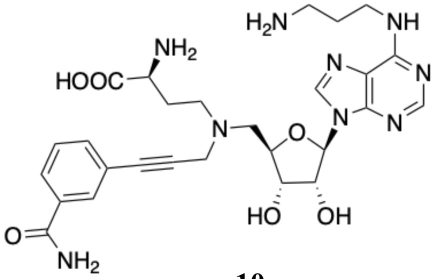
H2N
NH
NH2
HOOC
N
O
HO
OH
NH2
O

**10**

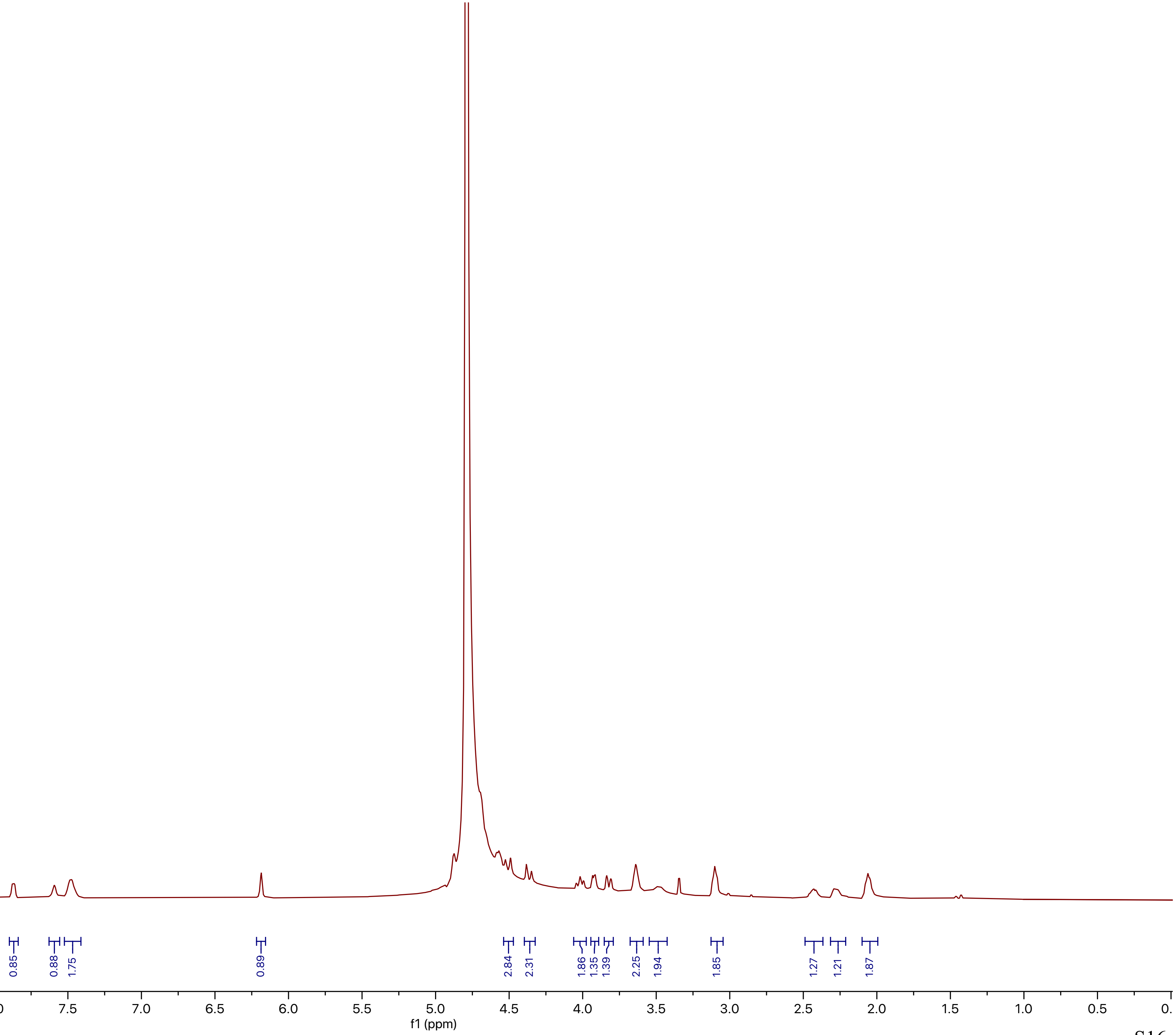
1.00
0.84
0.85
0.88
1.75
0.89
2.84
2.31
1.86
1.35
1.39
2.25
1.94
1.85
1.27
1.21
1.87
9.5
9.0
8.5
8.0
7.5
7.0
6.5
6.0
5.5
5.0
4.5
4.0
3.5
3.0
2.5
2.0
1.5
1.0
0.5
f1 (ppm)

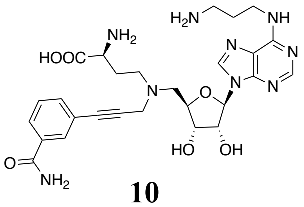


**10**

#### HRMS

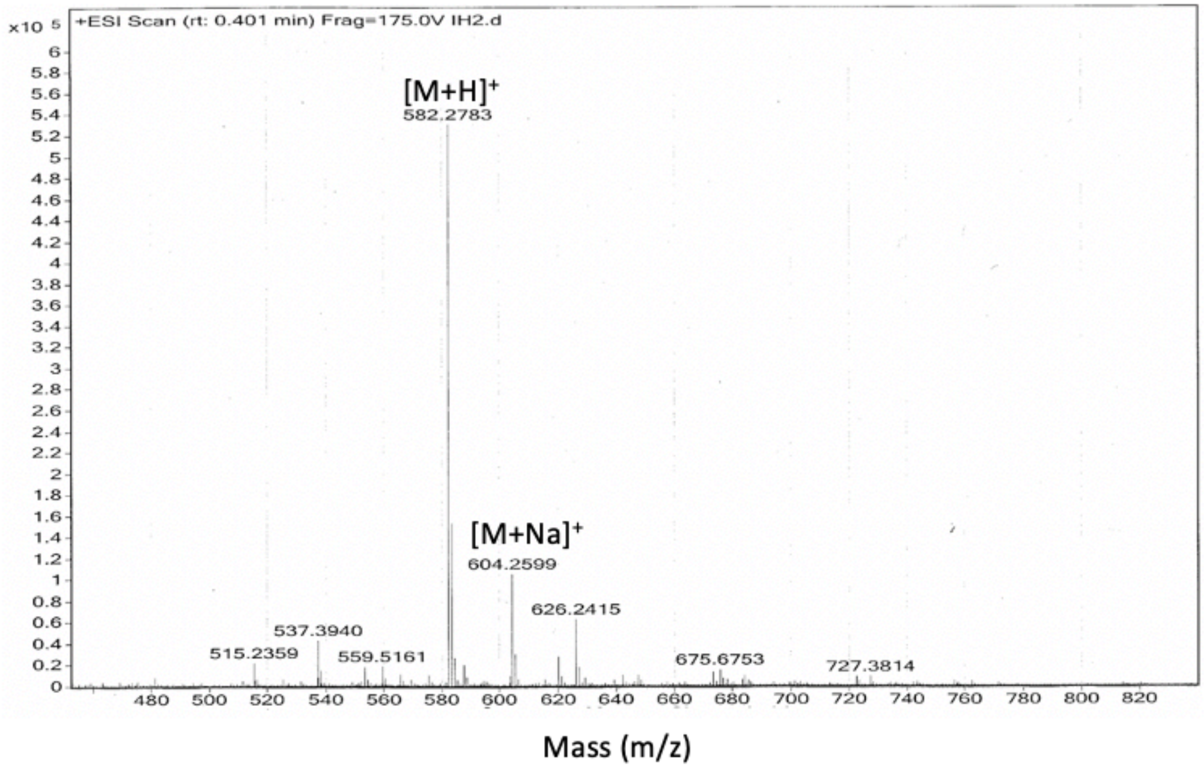


#### HPLC

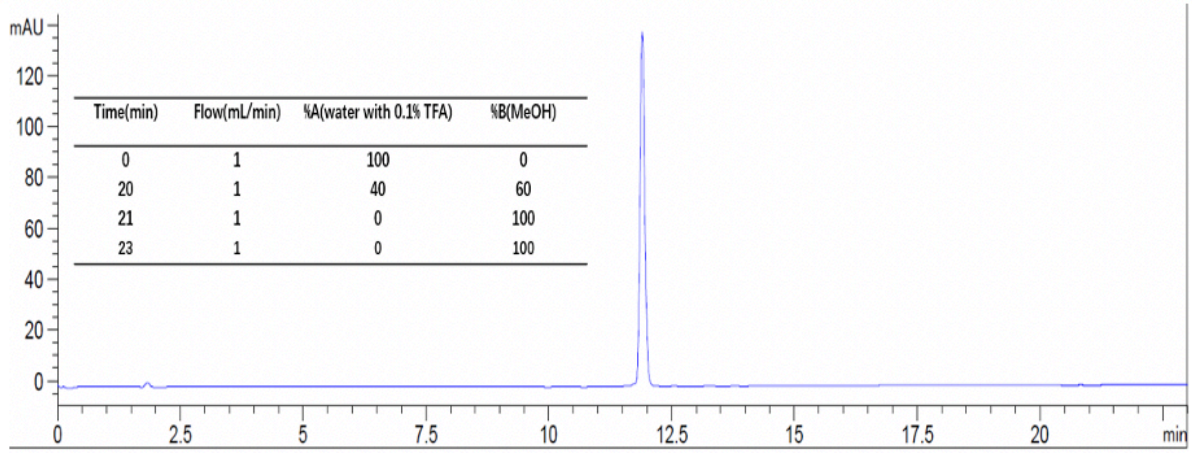


| Time(min) | Flow(mL/min) | %A(water with 0.1% TFA) | %B(MeOH) |
|---|---|---|---|
| 0 | 1 | 100 | 0 |
| 20 | 1 | 40 | 60 |
| 21 | 1 | 0 | 100 |
| 23 | 1 | 0 | 100 |

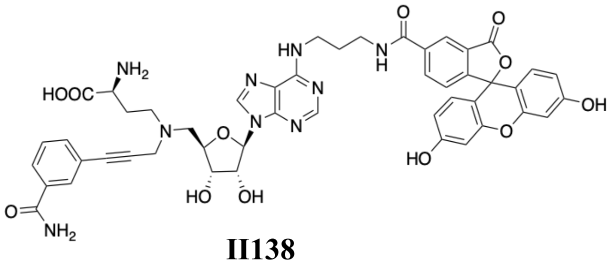

NH2
HOOC
N
HN
N
N
N
N
O
H
O
O
O
O
OH
HO
O
HO
OH
O
NH2


**II138**

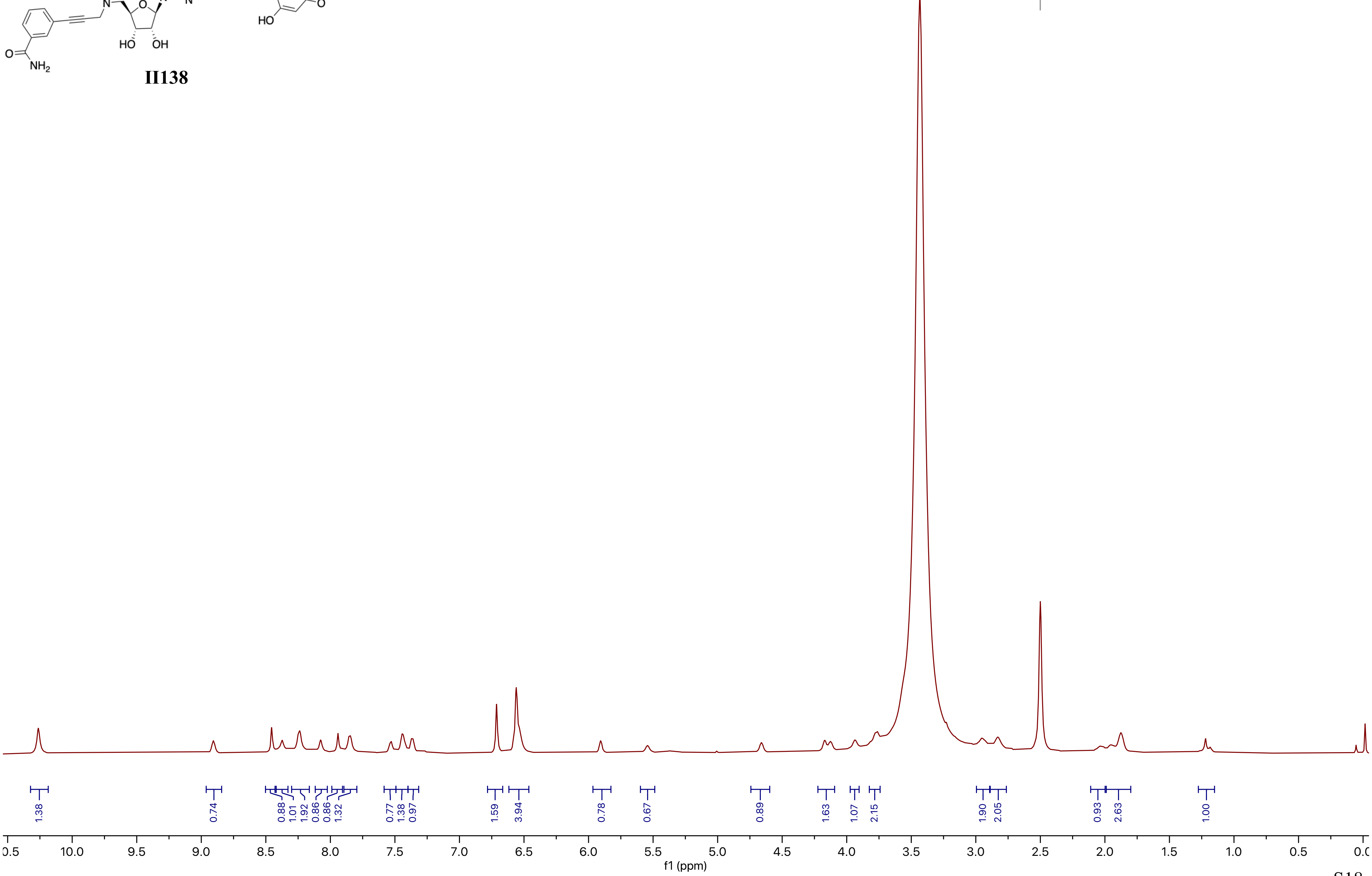

2.50 DMSO-d6
1.38
0.74
0.88
1.01
1.92
0.86
0.86
1.32
0.77
1.38
0.97
1.59
3.94
0.78
0.67
0.89
1.63
1.07
2.15
1.90
2.05
0.93
2.63
1.00
10.0
9.5
9.0
8.5
8.0
7.5
7.0
6.5
6.0
5.5
5.0
4.5
4.0
3.5
3.0
2.5
2.0
1.5
1.0
0.5
f1 (ppm)

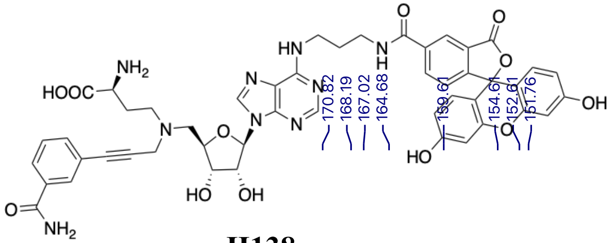

**II138**

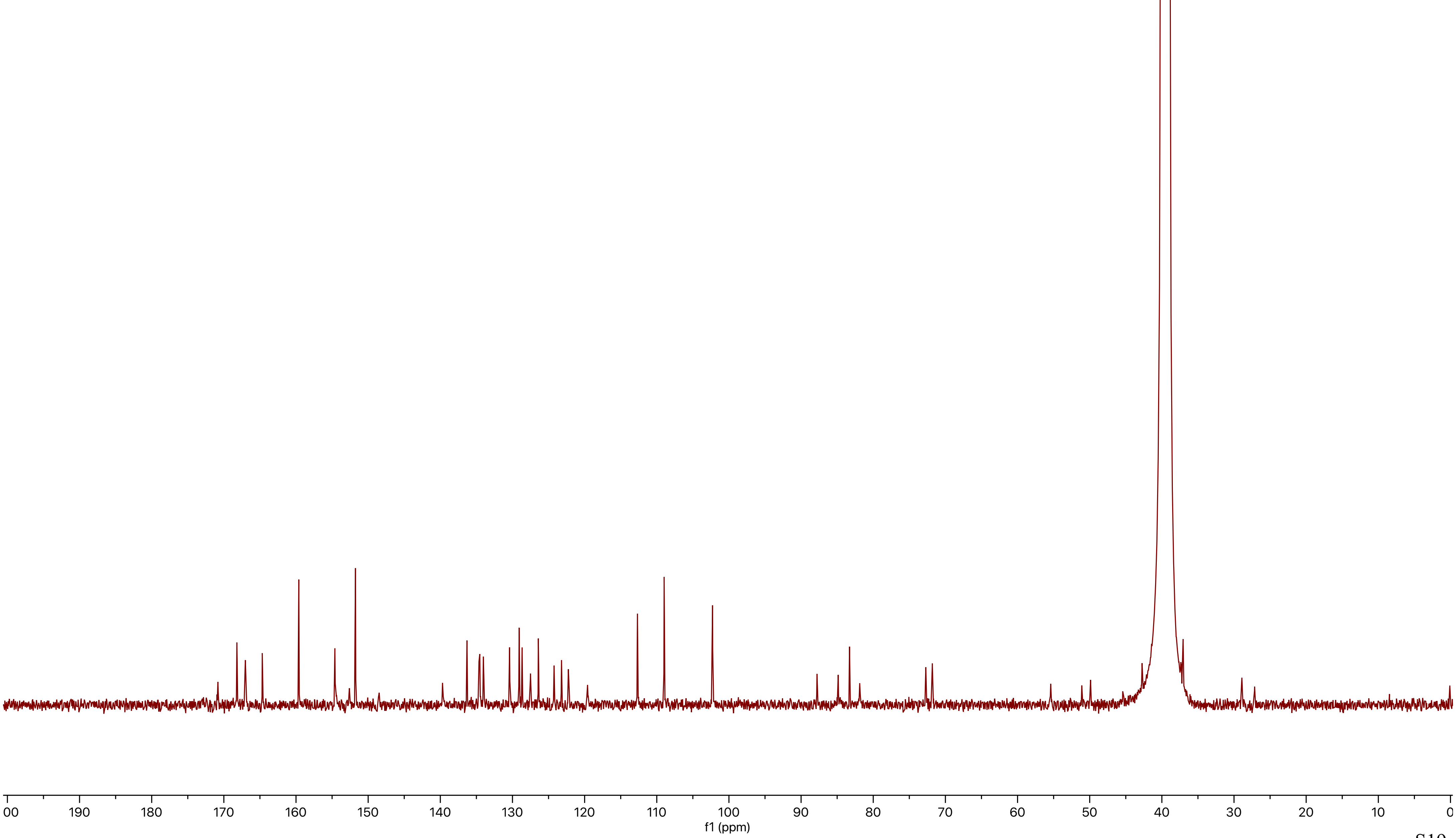
170.82
168.19
167.02
164.68
159.61
154.61
152.61
151.76
139.69
136.31
134.64
134.53
134.04
130.40
129.07
128.67
127.51
126.41
124.23
123.20
122.25
119.59
112.68
108.98
102.27
87.80
84.87
83.29
81.88
72.72
71.82
55.41
51.09
49.88
42.74
39.52 DMSO-d6
37.54
37.06
28.92
27.14
00
190
180
170
160
150
140
130
120
110
100
90
80
70
60
50
40
30
20
10
0
f1 (ppm)

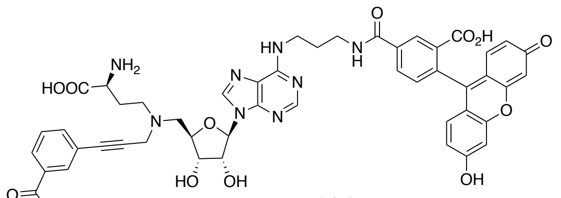


**II138**

### HRMS

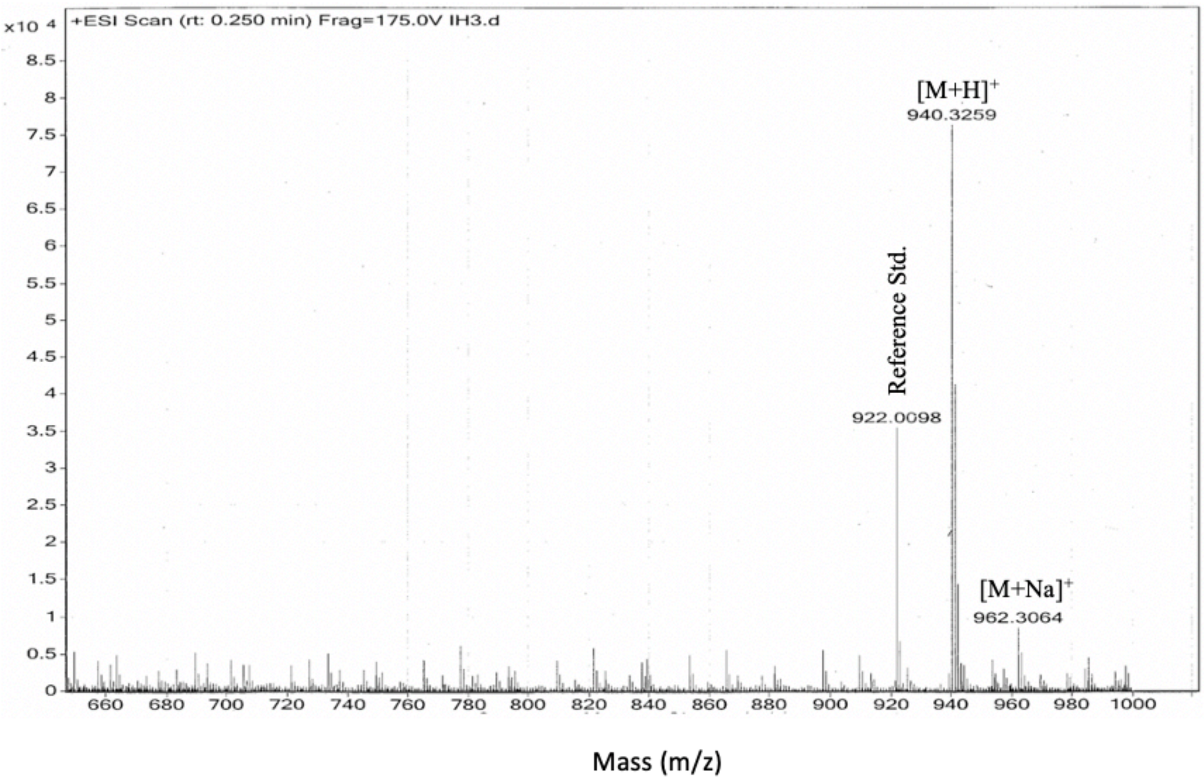


### HPLC

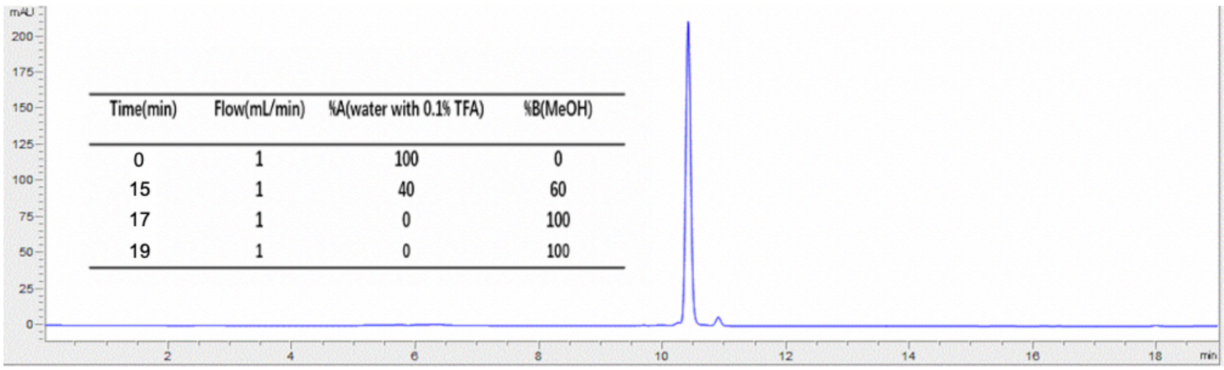

| Time(min) | Flow(mL/min) | %A(water with 0.1% TFA) | %B(MeOH) |
|---|---|---|---|
| 0 | 1 | 100 | 0 |
| 15 | 1 | 40 | 60 |
| 17 | 1 | 0 | 100 |
| 19 | 1 | 0 | 100 |
